# Glutathione-mediated S-Nitrosylation of WRKY75 modulates *PHT1;5* expression to orchestrate phosphate homeostasis in *Arabidopsis*

**DOI:** 10.1101/2022.11.03.515049

**Authors:** Ranjana Shee, Dibyendu Shee, Tanushree Saha, Salman Sahid, Soumitra Paul, Riddhi Datta

## Abstract

Phosphorus (P) is essential for plant growth, yet inorganic phosphate (Pi) is often limiting in soil. Plants respond to Pi limitation by activating P transporters and starvation genes, which boost Pi uptake or redistribute P. While glutathione (GSH) regulates iron and zinc homeostasis in Arabidopsis, its role in P homeostasis under Pi limitation remains elusive. Here, we show that GSH-S-nitrosoglutathione (GSNO) module regulates source-to-sink P allocation in Arabidopsis by transcriptionally activating the phloem-localised transporter *AtPHT1;5* via the transcription factor *At*WRKY75. GSH-deficient mutants, *cad2-1* and *pad2-1*, exhibited sensitivity to Pi-starvation with altered root system architecture, reduced anthocyanin accumulation, and disrupted root–to-shoot P transport. *AtPHT1;5* transcript levels were markedly reduced in these mutants but were positively responsive to exogenous GSH or GSNO. The *pht1;5* mutant failed to respond to GSH treatments, demonstrating that *AtPHT1;5* is essential for GSH-mediated P mobilisation. Pharmacological and genetic manipulation of nitric oxide (NO)/GSNO levels implicated GSNO as an essential intermediary. In depth molecular analyses identified *At*WRKY75 as a Pi- and GSH/GSNO-responsive activator that binds w-box motifs in the *AtPHT1;5* promoter. This was further supported by the lack of *AtPHT1;5* induction in the *wrky75* mutants. Finally, *At*WRKY75 is S-nitrosylated and stabilised by GSH/GSNO, providing a mechanistic link between redox/NO signalling and transcriptional control of P transport. We propose that elevated GSH under Pi limitation is converted to GSNO, which S-nitrosylates AtWRKY75 and promotes *AtPHT1;5* expression, thus driving source-to-sink P reallocation. This pathway integrates redox and NO signalling with transcriptional regulation to maintain P homeostasis under Pi starvation.

**Summary statement:** Glutathione-S-nitrosoglutathione module regulates phosphate translocation in Arabidopsis under phosphate starvation via transcriptional activation of *AtPHT1;5* gene by *At*WRKY75 transcription factor.

## Introduction

Phosphorus (P) is an essential macronutrient that regulates various physiological phenomena in plants. It serves as a crucial component of nucleic acids, phospholipids, and different sugar-phosphate intermediates. Moreover, the phosphorylation of regulatory proteins and enzymes plays a critical role in modulating various signal transduction pathways (Poirier and Bucher, 2002). In plants, inorganic phosphate (Pi) is the primary source of P taken up by the roots from the soil. However, Pi is poorly available to plants due to its high reactivity with soil calcium, aluminium, or iron (Fe), leading to its precipitation or incorporation into organic compounds (Bieleski, 1973). Therefore, Pi deficiency is a severe constraint on plant growth and development. Plants have evolved intricate mechanisms to improve Pi uptake and utilisation in the rhizosphere and to reallocate the internal Pi reserve to maintain P homeostasis under Pi-limited conditions. P deficiency alters root system architecture (RSA), impairs photosynthesis, triggers the synthesis of acid phosphatases (APases), ribonucleases, and organic acids, and induces anthocyanin and starch accumulation in plants (Vance et al., 2003). Since the concentration of Pi in a plant cell is 10^2^-10^3^ times higher than that in the soil, plants transport Pi against the concentration gradient with the help of different low and high-affinity P transporters (Shin et al., 2004). Earlier, three important P transporter families, *viz.,* the phosphate transporter (PHT) family, the SYG1/Pho81/XPR1 (SPX) domain-containing protein family, and the sulfate transporter (SULTR)-like family, were reported in plants (Liu et al., 2017). Among them, the PHT family transporters, which play a crucial role in Pi acquisition and P allocation, are categorised into five groups based on their subcellular localisation (Liu et al., 2016). For example, PHT1 localises on the plasma membrane, while PHT2 and PHT4 are on the chloroplast membrane. On the other hand, PHT3 and PHT5 localise on mitochondrial and vacuolar membranes, respectively.

The PHT1 family of P transporters comprises nine members in Arabidopsis, located on chromosomes 1, 2, 3, and 5 (Mudge et al., 2002). Interestingly, four PHT1 family genes, *PHT1;1*, *PHT1;2*, *PHT1;3*, and *PHT1;6*, are clustered on chromosome 5, indicating a gene duplication during the evolution of the PHT1 family in the Arabidopsis genome (Mudge et al., 2002). The family functions as H^+^/Pi symporters during Pi uptake from the soil and its distribution within plants (Zhang et al., 2014). The PHT1;1 plays a dominant role in Pi uptake from the rhizosphere under high P conditions, while PHT1;4 predominantly helps in Pi acquisition under P-limited conditions. Besides, PHT1;2, PHT1;3, and PHT1;5 also function in Pi uptake from soil (Shin et al., 2004; Nagarajan et al., 2011; Ayadi et al., 2015). Apart from Pi acquisition by roots, P allocation and transport to different tissues are also crucial for P homeostasis in plants. The PHT1;8 and PHT1;9 transporters function in root-to-shoot P translocation in co-operation with the PHO1, PHT1;3 and PHT1;4 (Mudge et al., 2002; Lapis-Gaza et al., 2014). Furthermore, P rescue from senescing leaves provides a means to recycle P reserves in plants. PHT1;5 is vital to the process. Similarly, the PHT1;5 transporter is implicated in P redistribution from source to sink tissues in response to developmental signals and soil P status (Nagarajan et al., 2011). Although Pi deprivation dramatically increases the expression of several *PHT1* genes, others are expressed irrespective of the soil Pi concentration. The low Pi signal is often transduced by several P-responsive transcription factors (TFs) like PHOSPHATE STARVATION RESPONSE 1 (PHR1), WRKY, and MYB domain-containing families to activate specific *PHT1* transporters (Rubio et al., 2001; Gu et al., 2016). Tight control of P transporters allows efficient Pi use in plants during P deficiency.

Glutathione (GSH) is a ubiquitous tripeptide that serves as a master regulator of plant growth, development, and stress response. It functions in reactive oxygen species (ROS) detoxification, redox homeostasis, and defence signalling network under various biotic and abiotic stress conditions (Foyer and Noctor, 2011; Noctor et al., 2012). In addition, GSH can confer tolerance to various forms of metal toxicity in plants by activating phytochelatin synthesis (Seth et al., 2012; Jozefczak et al., 2014). Several GSH-deficient Arabidopsis mutants with varying levels of susceptibility to biotic and abiotic stress factors have been identified. Among them, the *cadmium sensitive 2-1* (*cad2-1*) mutant displays severe sensitivity to cadmium and possesses around 30% GSH content than that in wild-type plants (Howden et al., 1995). On the other hand, the *phytoalexin-deficient2-1* (*pad2-1*) mutant, which contains 22% GSH, exhibits susceptibility to various pathogen infections (Glazebrook and Ausubel, 1994; Parisy et al., 2007).

GSH also modulates nutrient homeostasis in Arabidopsis. For example, the *zir1* mutant that contains 15% GSH displayed increased sensitivity to Fe-limited conditions (Shanmugam et al., 2012). During Fe deficiency, GSH maintains redox homeostasis in the cell and increases Fe uptake and accumulation by activating Fe-responsive genes such as *FRO2*, *FIT1*, and *IRT1* in Arabidopsis (Shanmugam et al., 2015). Earlier, we reported that the GSH-(S-nitrosoglutathione) GSNO module regulates subcellular Fe homeostasis by modulating NRAMP3, NRAMP4, and PIC1 transporters during Fe deficiency (Shee et al., 2022). However, the role of GSH in modulating the P transport or homeostasis remains unexplored. Although GSH-indole butyric acid (IBA) crosstalk was reported to alter the RSA under Pi-deprived conditions, the mechanism of GSH-mediated P homeostasis is not elucidated (Trujilo Hernandez et al., 2020). Here, we demonstrate that the GSH-GSNO module also regulates source-to-sink P transport via transcriptional activation of the *AtPHT1;5* transporter through the *At*WRKY75 TF under altered Pi conditions.

## Results

### GSH-deficient mutants display sensitivity to Pi deprivation

To determine whether GSH regulates P homeostasis, we analysed GSH-deficient mutants *cad2-1* and *pad2-1*, along with wild-type (Col-0) plants under Pi-deprived and Pi-sufficient conditions. The standard Murashige and Skoog (MS) medium (1.25 mM KH_2_PO_4_; +P) was Pi-sufficient, while the Pi-deficient medium (0 mM KH_2_PO_4_; 0.5 mM K_2_SO_4_; -P) (Dong et al., 2017) lacked Pi. Under Pi sufficiency, neither mutant showed any phenotypic differences compared with Col-0. After 7 d of Pi starvation, all genotypes developed smaller rosettes, shorter primary roots, and increased lateral root density than under Pi sufficiency. However, *pad2-1* was more sensitive to Pi deficiency, with reduced rosette diameter, fewer lateral roots, and lower lateral root density than Col-0 (Fig. 1A-E). Chlorophyll content dropped in all genotypes during Pi deficiency, with the sharpest drop in *pad2-1* (Fig. 1F). Conversely, anthocyanin increase under Pi deprivation was significantly lower in *cad2-1* and *pad2-1* than Col-0 (Fig. 1G).

**FIGURE 1.**
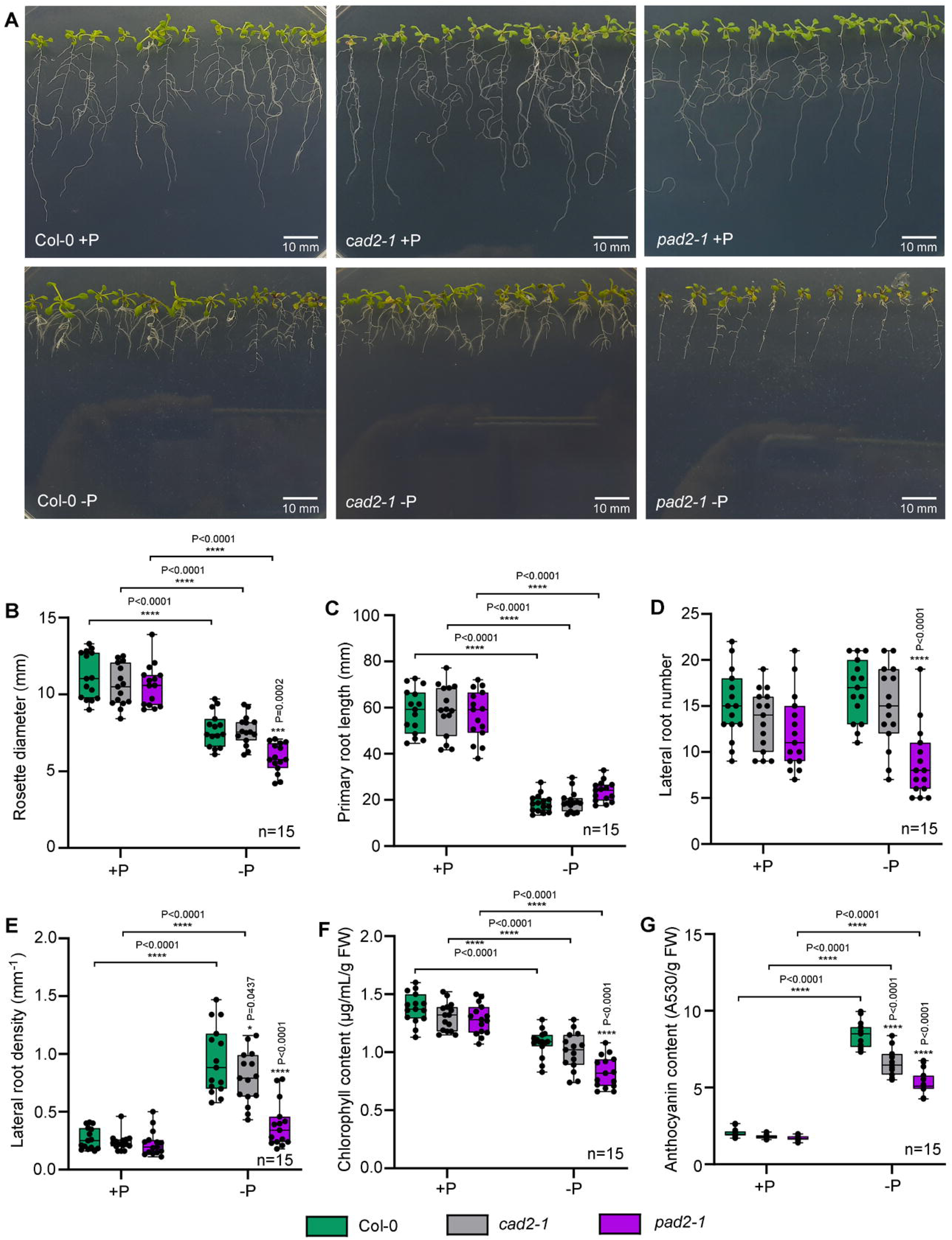
Response of GSH-deficient mutants to Pi starvation. 7 d old MS grown seedlings of Col-0, *cad2-1* and *pad2-1*were exposed to +P and -P conditions for 7 d and analyzed. (A) Representative images of plants grown under +P and –P conditions, (B) rosette diameter, (C) primary root length, (D) lateral root number, (E) lateral root density, (F) chlorophyll content, and (G) anthocyanin content. The experiment was independently repeated thrice and results were represented as mean±SEM (n=15). Statistical differences between the genotypes and between the Pi-sufficient (+P) and Pi deficient (-P) conditions were analyzed by two-way ANOVA followed by Dunnett’s multiple comparison test and Tukey’s multiple comparison test respectively. Statistical significances were denoted by asterisks in the respective panels.

To assess potential P-GSH interactions, we measured GSH, Pi, and total P content under Pi-sufficient and Pi-deficient conditions. Under Pi-sufficient conditions, root Pi and total P contents were significantly lower in both *cad2-1* and *pad2-1* mutants. In response to Pi deprivation, however, *pad2-1* roots accumulated more Pi and total P than *cad2-1* and Col-0 (Fig. 2A, B). Also, total root GSH increased in Col-0, while the GSH:GSSG ratio decreased in all three genotypes under Pi starvation (Fig. 2C-D). In shoots, total GSH increased in Col-0, and the GSH:GSSG ratio decreased in all three genotypes under Pi limitation, whereas Pi and total P content showed contrasting patterns (Fig. 2E-H). Shoot Pi and total P were higher in mutants under Pi-sufficient conditions, but lower under Pi deficiency.

**FIGURE 2.**
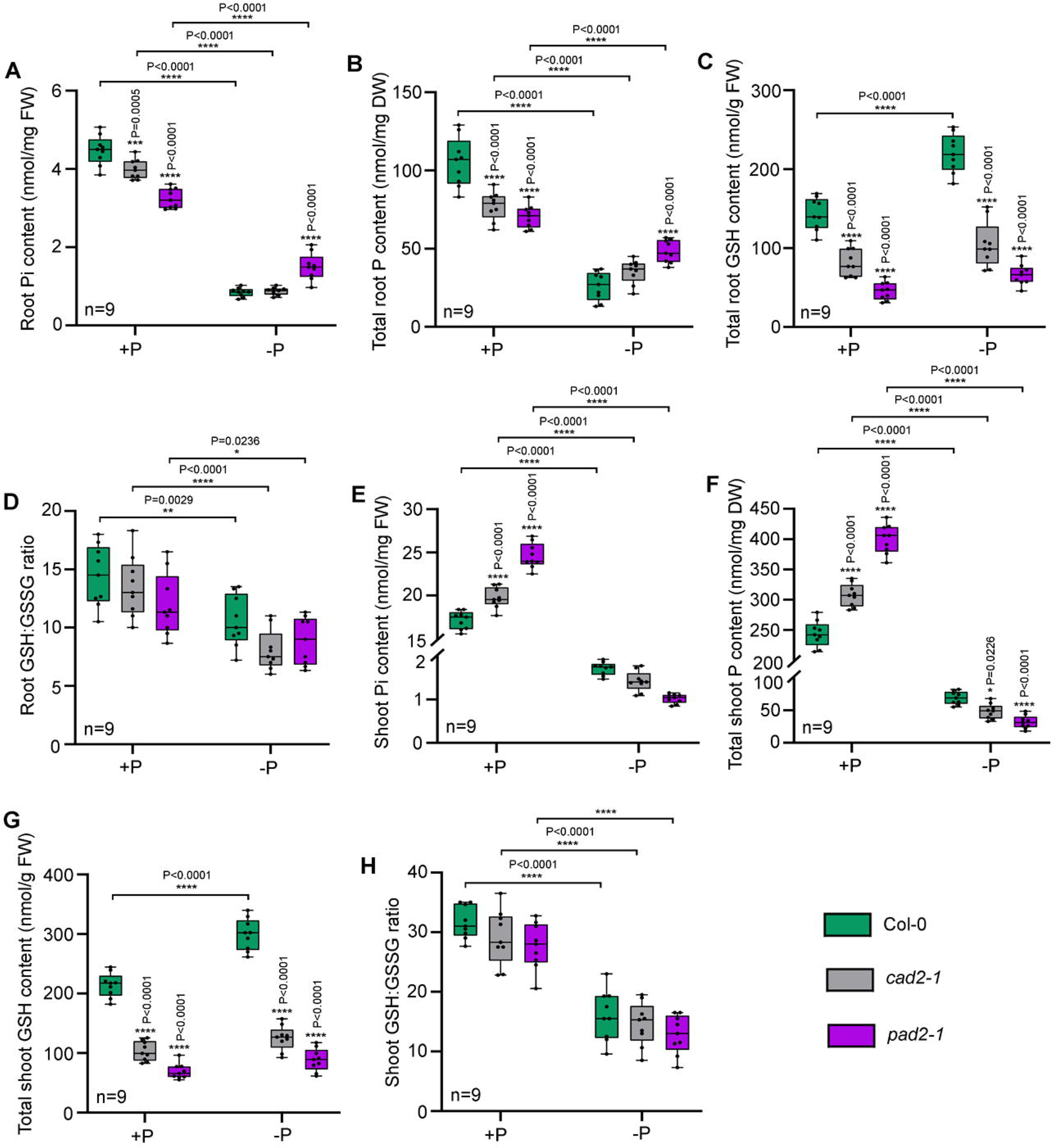
Estimation of Pi and total P contents, total GSH and GSH:GSSG ratio under +P and -P conditions. 7 d old MS grown Col-0, *cad2-1* and *pad2-1*seedlings were exposed to -P condition for 7 d and analyzed. Control plants were maintained in the MS medium for the entire period. (A) Root Pi content, (B) total root P content (C) total root GSH content, (D) root GSH:GSSG ratio, (E) shoot Pi content, (F) total shoot P content, (G) total shoot GSH content, and (H) shoot GSH:GSSG ratio. The experiment was independently repeated thrice and results were represented as mean±SEM (n=9). Statistical differences between the genotypes and between the Pi-sufficient (+P) and Pi deficient (-P) conditions were analyzed by two-way ANOVA followed by Dunnett’s multiple comparison test and Tukey’s multiple comparison test respectively. Statistical significances were denoted by asterisks in the respective panels.

### GSH regulates AtPHT1;5 to modulate P transport

The sensitivity of the GSH-deficient mutants to Pi starvation and their altered P status prompted us to investigate whether GSH regulates P transporter genes. To this end, we analysed the expression of several known P transporters in the root and shoot under Pi-sufficient conditions and compared their relative transcript abundance between Col-0 and the GSH-deficient mutant. Interestingly, the *AtPHT1;5* expression was significantly down-regulated in both the mutants compared to the Col-0 plants (Fig. 3A-B). Although Pi deficiency induced the *AtPHT1;5* expression in all three genotypes, its transcript abundance was significantly lower in the mutants than in the Col-0 plants (Fig. 3C-D).

**FIGURE 3.**
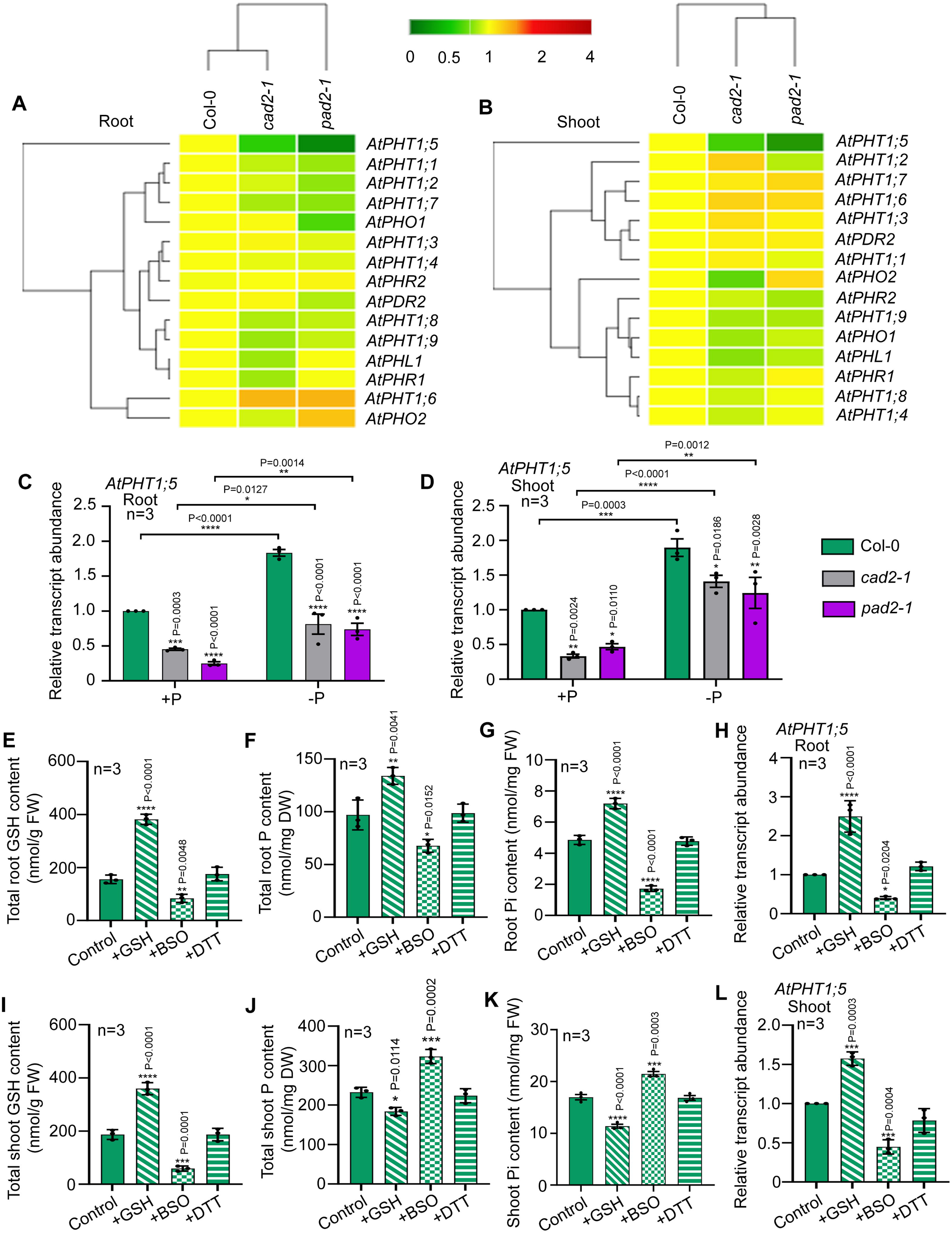
Expression analyses of different P transporters under altered GSH conditions. 14 d old MS grown Col-0, *cad2-1*, and *pad2-1* plants were analyzed for the expression pattern of various P transporter genes. Expression of different P transporters from root (A) and shoot (B) tissues was depicted as heat map. Relative transcript abundance of *AtPHT1;5* gene was analyzed from root (C) and shoot (D) of 7 d old Col-0, *cad2-1*, and *pad2-1*seedlings grown under +P and -P conditions for additional 7 d. 14 d old MS grown Col-0 plants were treated with 100 μM GSH for 72 h, or 1 mM BSO for 72 h, or 5 mM DTT solutions for 24 h and analyzed. Control plants were maintained in half strength MS medium for the entire duration. (E) Total root GSH content, (F)total root P content,(G) root Pi content, (H) relative transcript abundance of *AtPHT1;5*gene in roots, (I) total shoot GSH content, (J) total shoot P content, (K) shoot Pi content and (L) relative transcript abundance of *AtPHT1;5*gene in shoots. The experiment was independently repeated thrice and results were represented as mean±SEM (n=3). Statistical differences between the genotypes or treatments were analyzed by one-way ANOVA (A,B, E-L) or two-way ANOVA (C-D) followed by Dunnett’s multiple comparison test and statistical significances were denoted by asterisks in the respective panels.

Further, we examined the effect of altered GSH on *AtPHT1;5* expression in the Col-0 plants under Pi-sufficient and deficient conditions. The total GSH content increased in response to exogenous GSH treatment, while the GSH biosynthesis inhibitor, buthionine sulphoximine (BSO), decreased it. However, treatment with the non-specific reducing agent dithiothreitol (DTT) did not alter total GSH content (Fig. 3E, I). Interestingly, the root Pi and total P content increased significantly in response to GSH treatment but decreased in response to BSO treatment (Fig. 3F, G). The shoot, on the other hand, displayed an opposite pattern (Fig.3J, K). However, DTT treatment failed to alter P and Pi contents in either the root or the shoot. In line with this observation, the *AtPHT1;5* expression was up-regulated in response to GSH treatment and down-regulated by BSO treatment, while DTT did not alter its expression in root as well as shoot (Fig. 3H, L). A similar pattern was observed under Pi-deficient conditions as well (Fig S1). Further, we treated the GSH-deficient *pad2-1* mutant with exogenous GSH and DTT. While total GSH content increased with GSH treatment, no significant alteration was observed with DTT treatment. In response to exogenous GSH, the Pi and total P contents increased in roots but decreased in shoots, indicating a role of GSH in modulating P homeostasis (Fig S2). In addition, *AtPHT1;5* expression was induced by GSH treatment, whereas DTT treatment did not alter its transcript abundance (Fig S2). Together, these observations strongly suggest that GSH mediates regulation of P transport through the *AtPHT1;5* transporter.

### GSH modulates P mobilization during Pi starvation by regulating AtPHT1;5 expression

To further examine the role of GSH in P mobilisation, we analysed the response of *pht1;5* mutant to exogenous GSH. We generated complementation (*pht1;5/pPHT1;5::PHT1;5*) lines by introducing the *AtPHT1;5* gene, under control of its native promoter, into the *pht1;5* mutant background and analyzed their response (Fig. 4A). To check the correct complementation of the *pht1;5* mutant, we analysed the expression of the *AtPHT1;5* gene in the WT, mutant and the complementation lines. The expression levels in the WT and the complementation lines were found to be similar, while no expression was detected in the mutant line (Fig S3). In addition, the expression of the *AtPHT1;4* gene, a marker for Pi starvation, was analysed under Pi-sufficient and Pi-deficient conditions. Its expression was significantly induced by Pi starvation in all three genotypes (Fig. S4). Col-0, *pht1;5* and complementation lines had similar total GSH contents, which increased under Pi starvation. Exogenous GSH increased total GSH in all genotypes under both Pi-sufficient and Pi-deficient conditions (Fig. 4B, E). The *pht1;5* mutant showed altered root-to-shoot P allocation in both nutrient conditions. Under Pi sufficiency, it accumulated less Pi and P in roots but more in shoots compared to Col-0. In Pi-deficient conditions, Pi and P levels were higher in roots and lower in shoots. The complementation lines resembled Col-0 plants (Fig. 4C, D, F, G). Exogenous GSH enhanced root Pi and P in Col-0 and complementation lines under Pi-sufficiency, but reduced them during Pi starvation. In shoots, GSH reduced Pi and P under Pi-sufficiency but increased them under Pi deficiency. In contrast, exogenous GSH had no effect on *pht1;5* mutant root or shoot Pi and P content under either condition (Fig. 4C, D, F, G). *AtPHT1;5* expression was highly induced in GSH-treated and reduced in BSO-treated Col-0 plants. Complementation lines in altered GSH conditions showed expression patterns similar to Col-0 (Fig S3). This suggests GSH modulates root-to-shoot P allocation through *At*PHT1;5 to regulate P homeostasis during Pi starvation.

**FIGURE 4.**
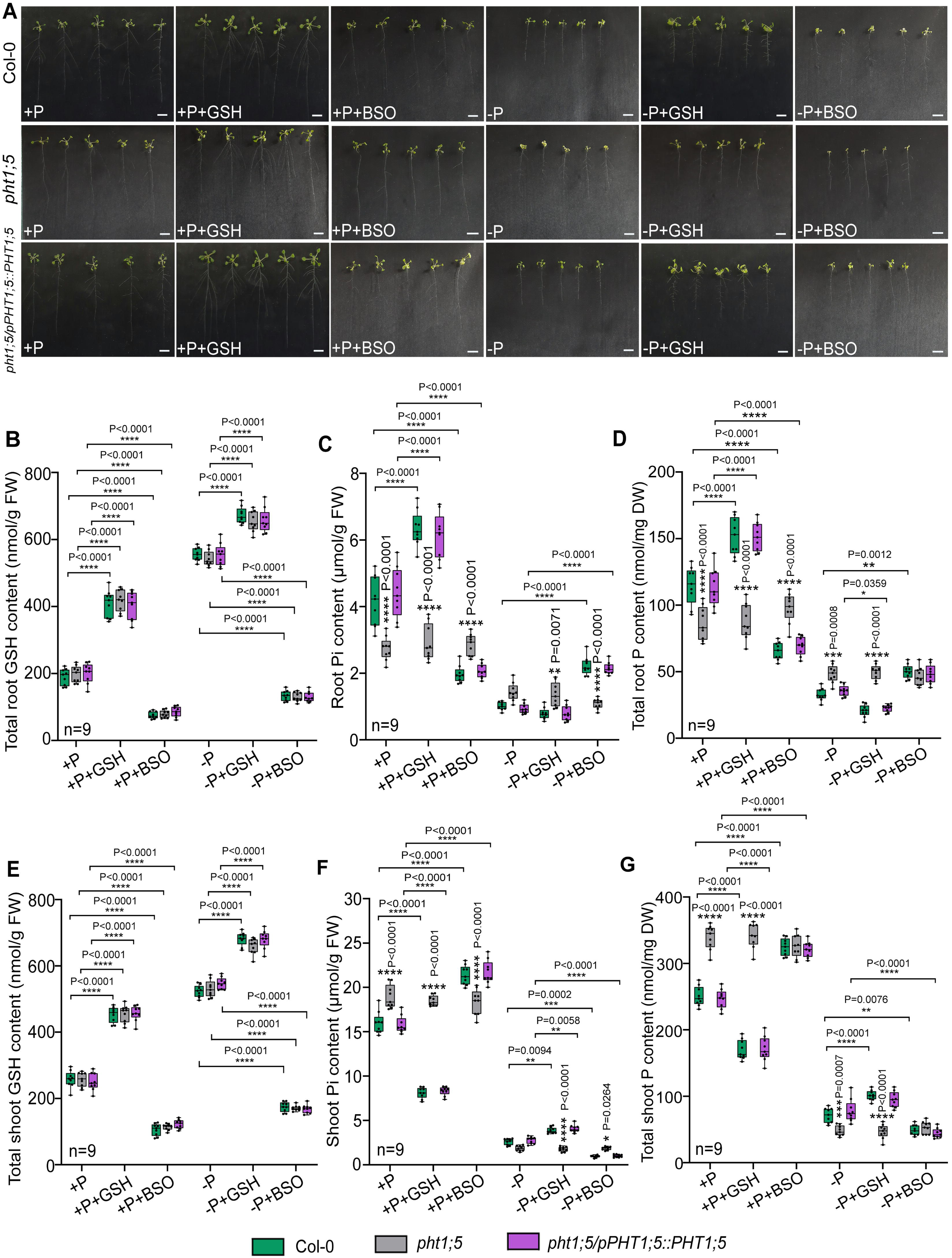
Response of *pht1;5* mutant and complementation lines to exogenous GSH and BSO under Pi-sufficient and -deficient conditions. 7 d old MS grown seedlings of Col-0, *pht1;5* mutant and complementation (*pht1;5/pPHT1;5::PHT1;5*) lines were exposed to +P and -P conditions for 7 d and treated with 100 μM GSH, or 1 mM BSO for 72 h and analyzed. (A) Representative images of plants grown under +P, +P+GSH,+P+BSO, –P, – P+GSH and –P+BSO conditions, (B) total root GSH content, (C) root Pi content (D) total root P content, (E) total shoot GSH content, (F) shoot Pi content and (G) total shoot P content. The experiment was independently repeated thrice and results were represented as mean±SEM (n=9). Statistical differences between the genotypes and between the conditions (+P, +P+GSH, +P+BSO, -P, –P+GSH and –P+BSO) were analyzed by two-way ANOVA followed by Dunnett’s multiple comparison test and Tukey’s multiple comparision test respectively. Statistical significances were denoted by asterisks in the respective panels. Scale bar: 10 mm

### GSNO Contributes to GSH-Mediated P Homeostasis

Since the GSH-GSNO interactions regulate multiple physiological processes in plants, we investigated whether GSNO is necessary for GSH-mediated P homeostasis. To explore this, we analysed the effects of GSH, GSNO, the NO donor SNP, the GSH inhibitor BSO, and the NO scavenger cPTIO on Col-0 plants grown under Pi-deficient conditions. The root Pi and P contents dropped significantly in response to Pi deficiency compared to the Pi-sufficient condition. Notably, when GSH and cPTIO were combined, or SNP and BSO were applied together under Pi-deficient conditions, root Pi and P contents increased relative to Pi deficiency alone (Fig. 5A, B). In the shoot, however, the Pi and P contents showed opposite patterns (Fig. 5D, E). To further examine the molecular response, qRT-PCR analysis revealed strong *AtPHT1;5* induction following GSH, GSNO, or SNP treatment under Pi deficiency, compared with Pi deficiency alone. However, co-treatment with BSO and SNP blocked this induction, mirroring the effect of Pi deficiency alone. Similarly, adding GSH and cPTIO together did not increase *AtPHT1;5* expression (Fig. 5C,F).

**FIGURE 5.**
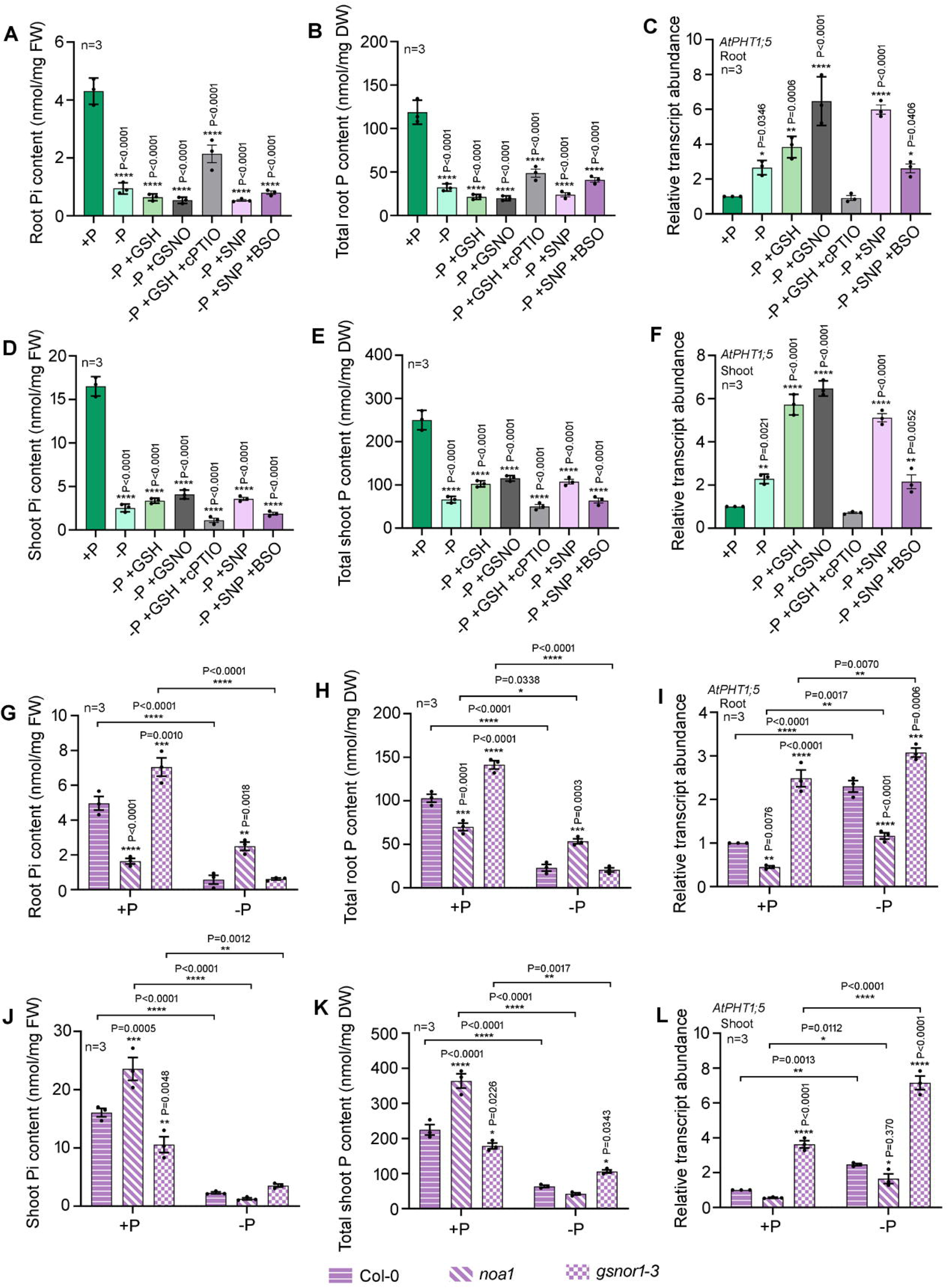
Response of Col-0 seedlings to exogenous GSH, GSNO, cPTIO, and SNP under Pi- deficient condition and responses of NO and GSNO mutant seedlings in response to Pi-starvation. 7 d old MS grown seedlings of Col-0 plants were exposed to -P conditions for 7 d and treated with 100 μM GSH, or 250 μM GSNO for 72 h, or 1 μM SNP for 48 h, or in combination of 100 μM GSH (72 h) and 0.5mM cPTIO (48 h), or in combination of 1 μM SNP (48 h) and 1 mM BSO (for 72 h) and analyzed. Control plants were maintained in half strength MS medium for the entire duration (A) root Pi content (B) total root P content (C) relative transcript abundance of *AtPHT1;5*gene in roots (D) shoot Pi content (E) total shoot P content (F) relative transcript abundance of *AtPHT1;5*gene in shoots. 7 d old MS grown seedlings of Col-0, noa*1* and *gsnor 1-3* were exposed to +P and -P conditions for 7 d and analyzed. (G) root Pi content, (H) total root P content (I) relative transcript abundance of *AtPHT1;5*gene from roots harvested from the mutants and WT seedlings grown in +P and –P condition (J) shoot Pi content, (K) total shoot P content (L) relative transcript abundance of *AtPHT1;5*gene from shoots harvested from the mutants and WT seedlings grown in +P and –P condition. The experiment was independently repeated thrice and results were represented as mean±SEM (n=3). Statistical differences between the conditions and between the genotypes (G-L) were analyzed by two-way ANOVA followed by Tukey’s multiple comparision test and Dunnett’s multiple comparison test respectively. Statistical significances were denoted by asterisks in the respective panels.

To build on these findings, we next measured Pi and P contents, as well as *AtPHT1;5* expression, in NO-related mutants *noa1* and *gsnor1-3*. Both Pi and P contents were significantly lower in the NO-deficient *noa1* roots, and higher in the shoots, while the GSNO-hyperaccumulating *gsnor1-3* mutant displayed higher levels in the roots and lower accumulation in the shoots under Pi-sufficient conditions. Under Pi deficiency, a reverse pattern was observed (Fig. 5G,H, J, K). Transcript levels were significantly downregulated in the *noa1* mutant but upregulated in the *gsnor1-3* mutant (Fig. 5I,L).

### GSH-GSNO module regulates transcriptional activation of the AtPHT1;5 gene

Since the GSH-GSNO module can regulate P allocation via the *AtPHT1;5* transporter, the next pertinent question was how they modulate the expression of this gene. To address this, we cloned the *AtPHT1;5* promoter (*pPHT1;5*) and generated transgenic Arabidopsis lines harbouring the *pPHT1;5*::*GUS* construct in the Col-0 background, along with a transgenic line containing the *CaMV35S::GUS* construct as a negative control. We then performed a histochemical GUS assay to analyse promoter activity under Pi-deficient conditions and after GSH and GSNO treatments. The results showed that *AtPHT1;5* promoter activity was strongly induced in Pi-deficient conditions, indicating a Pi-responsive nature (Fig. 6A). Exogenous GSH and GSNO significantly increased promoter activity, while DTT treatment had no effect. In contrast, BSO treatment decreased promoter activity, suggesting regulation by the GSH-GSNO module. Similarly, *uidA* gene expression was up-regulated in Pi-deficient conditions and with exogenous GSH and GSNO applications, but remained unchanged with DTT treatment, and was down-regulated with BSO treatment (Fig. 6B). Notably, neither Pi-deficiency nor other treatments altered GUS activity or *uidA* gene expression in the negative control line (Fig. 6C-D). Together, these findings demonstrate that the GSH-GSNO module regulates *AtPHT1;5* expression at the transcriptional level.

**FIGURE 6.**
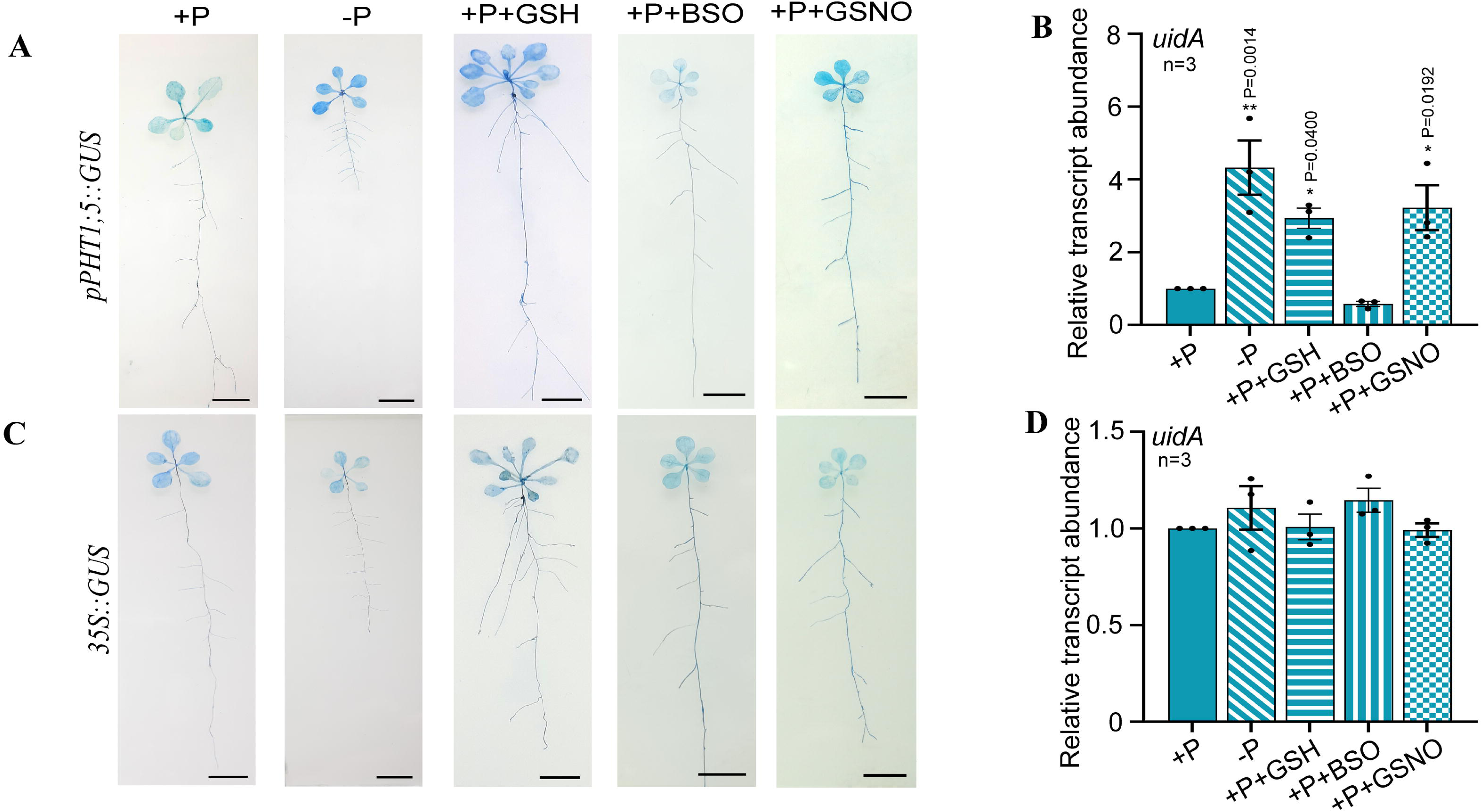
Promoter analysis of the *AtPHT1;5* gene. The seedlings of transgenic lines harboring *pPHT1;5::GUS* construct and the vector control (*35S::GUS*) lines were exposed to +P, –P, +P+GSH, +P+BSO, +P+GSNO conditions and used for promoter analysis by histochemical GUS assay and qRT-PCR analysis for *GUS (uidA)* gene expression. (A) Representative images of histochemical GUS staining of transgenic line harboring*pPHT1;5::GUS* construct, (B) relative transcript abundance of *GUS* gene of the same, (C) representative images of histochemical GUS staining of vector control line harboring the *35S::GUS* construct(negative control), and(D) relative transcript abundance of *GUS* gene of the same. The experiment was independently repeated thrice. For relative transcript abundance, results were represented as mean±SEM (n=3). Statistical differences between the treatments were analyzed by one-way ANOVA followed by Dunnett’s multiple comparison test and statistical significance of relative gene expression was denoted in the respective panels. Scale bar: 5 mm

### GSH-GSNO module regulates AtPHT1;5 expression via AtWRKY75 TF

GSH cannot directly bind to the promoter to regulate transcription. Therefore, we were curious whether any TFs are involved in the process. Several WRKY and MYB TFs, like *At*WRKY6, *At*WRKY42, *At*WRKY45, *At*WRKY75, *At*MYB2, and *At*MYB62, are known to regulate Pi-responsive genes in Arabidopsis. We performed a yeast one-hybrid (Y1H) assay to determine whether these TFs interact with the *AtPHT1;5* promoter and regulate its transcription. We observed that the *At*WRKY42, *At*WRKY75, and *At*MYB2 could bind to the *AtPHT1;5* promoter, while others did not (Fig. 7A). Next, we wondered if all three of these TFs were involved in the GSH-mediated regulation of *AtPHT1;5* under Pi-deficient conditions. Interestingly, we found that among these TFs, only the *AtWRKY75* expression was down-regulated in the seedlings of GSH-deficient mutants, *cad2-1*, and *pad2-1* (Fig. 7B). We further analysed the relative transcript abundance of the *AtPHT1;5* gene in the three TF mutant seedlings under Pi-sufficient and Pi-deficient conditions. Supporting our previous observation, the *AtPHT1;5* expression was induced in response to Pi starvation in the Col-0, *wrky42*, and *myb2* mutants but not in the *wrky75* mutant (Fig. 7C). This observation strongly indicated that the *At*WRKY75 TF is essential for the Pi-responsiveness of the *AtPHT1;5* gene. The dual luciferase transactivation assay further confirmed *AtPHT1;5* promoter activation by *At*WRKY75 (Fig. 7D). We next aimed to identify the region in the *AtPHT1;5* promoter that *At*WRKY75 specifically binds. *In silico* analysis revealed the presence of three w-box (TTGAC[CT]) motifs in the *AtPHT1;5* promoter region at -99, -176, and -201 positions (Fig. S5). Chromatin immunoprecipitation (ChIP)-qPCR assay confirmed the binding of the *At*WRKY75 TF to the w-box regions of the *AtPHT1;5* promoter with an enhanced binding affinity in response to Pi deficiency, exogenous GSH or GSNO treatment (Fig. 7E). Next, we wanted to analyse the *AtPHT1;5* promoter activity when the *At*WRKY75 TF was absent. Therefore, we expressed the *pPHT1,5::GUS* construct in the *wrky75* mutant background. A *CaMV35S::GUS* construct in the same background served as a negative control. We observed that neither Pi starvation nor exogenous GSH or GSNO treatment induced *AtPHT1;5* promoter activity, as evidenced by histochemical GUS activity (Fig. 7F). Together, these observations strongly suggest that transcriptional induction of *AtPHT1;5* by the GSH-GSNO module is mediated by AtWRKY75.

**FIGURE 7.**
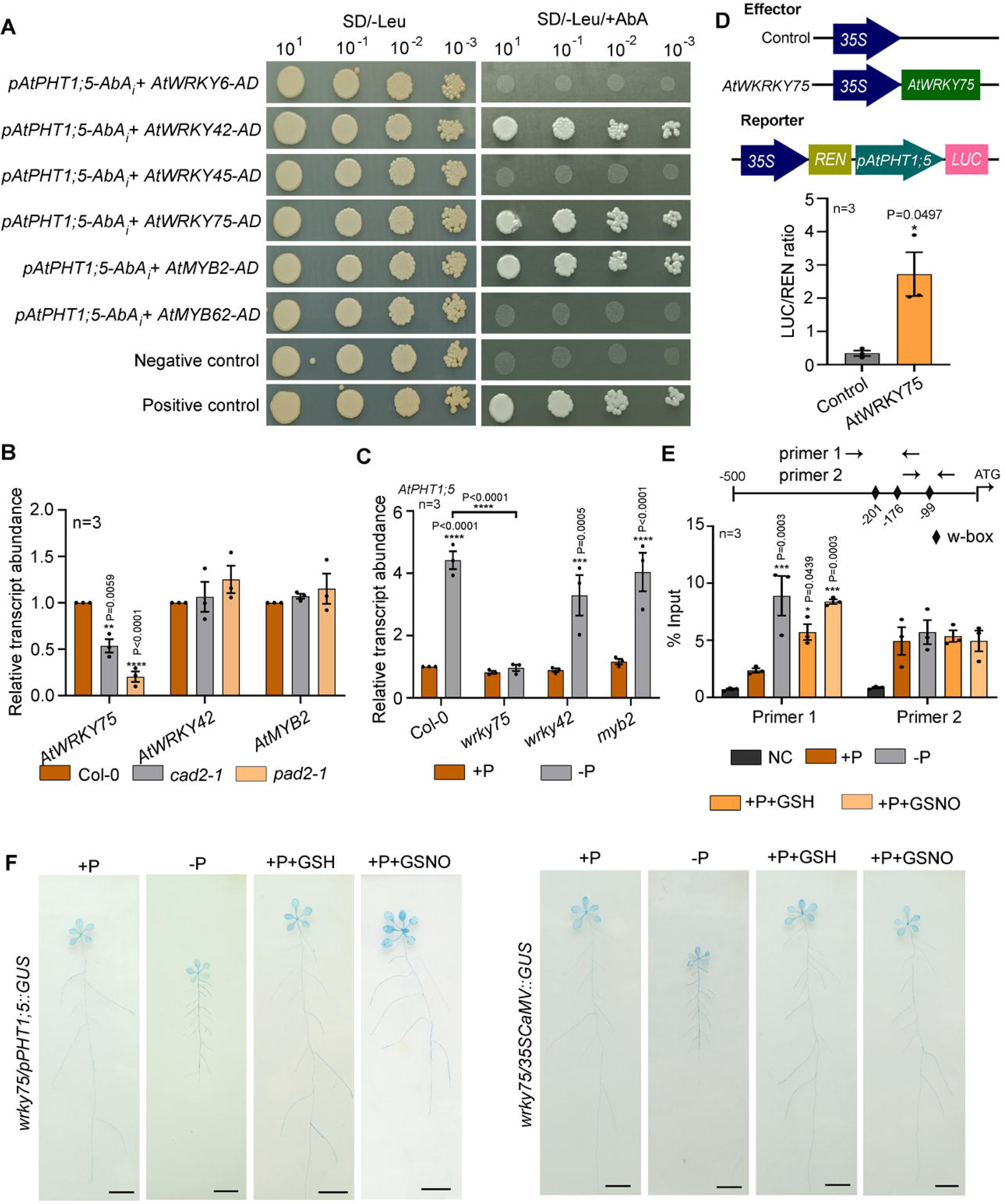
Identification of TFs interacting with *AtPHT1;5* promoter and its regulation by GSH and GSNO. (A) Y1H analysis for interaction of the P responsive TFs, *At*WRKY6, *At*WRKY42, *AtWRKY45*, *At*WRKY75, *At*MYB2, and *At*MYB62 with the *AtPHT1;5* promoter. (B) Relative transcript abundance of *AtMYB2*, *AtWRKY42* and *AtWRKY75* genes in the Col-0, *cad2-1* and *pad2-1* plants, (C) relative transcript abundance of *AtPHT1;5* gene in theCol-0, *wrky42*, *wrky75*, and *myb2* mutants grown under +P and –P condition. (D) Dual luciferase transctivation assay for *At*WRKY75-*AtPHT1;5* promoter. (E) ChIP-qPCR analysis for binding of *At*WRKY75 to the *AtPHT1;5* promoter under +P, -P, +P+GSH and +P+GSNO conditions. The position of the w-box motifs along with primer positions are marked on the promoter. NC: negative control. (F) Promoter analysis of the *AtPHT1;5* gene in the*wrky75* mutant background. Representative images of histochemical GUS staining of the transgenic *wrky75/pPHT1;5::GUS* and *wrky75/35S::GUS* (vector control) lines grown under +P, -P, +P+GSH and +P+GSNO conditions. The experiment was independently repeated thrice and results were represented as mean±SEM (n=3). Statistical differences between the treatments (B-C, E, G) were analyzed by one-way ANOVA followed by Dunnett’s multiple comparison test or student t test (D). Statistical differences between the genotypes (C) were carried out by Tukey’s multiple comparison tests and statistical significance was denoted in the respective panels.

### S-nitrosylation enhances the stability of AtWRKY75 TF

Since the GSH-GSNO module was found to induce *AtPHT1;5* promoter activity, we examined whether the *At*WRKY75 TF is S-nitrosylated. The protein sequence was scanned with GPS-SNO 1.0, which identified a single putative S-nitrosylation site (Fig. 8A, S6). Moreover, the DAN assay was performed to measure DAN□NAT conversion via NO released from the S□nitrosylated thiol group in proteins, and the results were compared with those for GSNO, which has a single S□nitrosylated Cys residue. The result validated S-nitrosylation of the *At*WRKY75 protein (Fig. 8B). Interestingly, the stability of the *At*WRKY75 protein was increased in response to GSH and GSNO treatment, further supporting our *in silico* data (Fig. 8C). We analysed the *At*PHT1;5 protein sequence as well, but no putative S-nitrosylation site could be predicted with sufficient confidence score.

**FIGURE 8.**
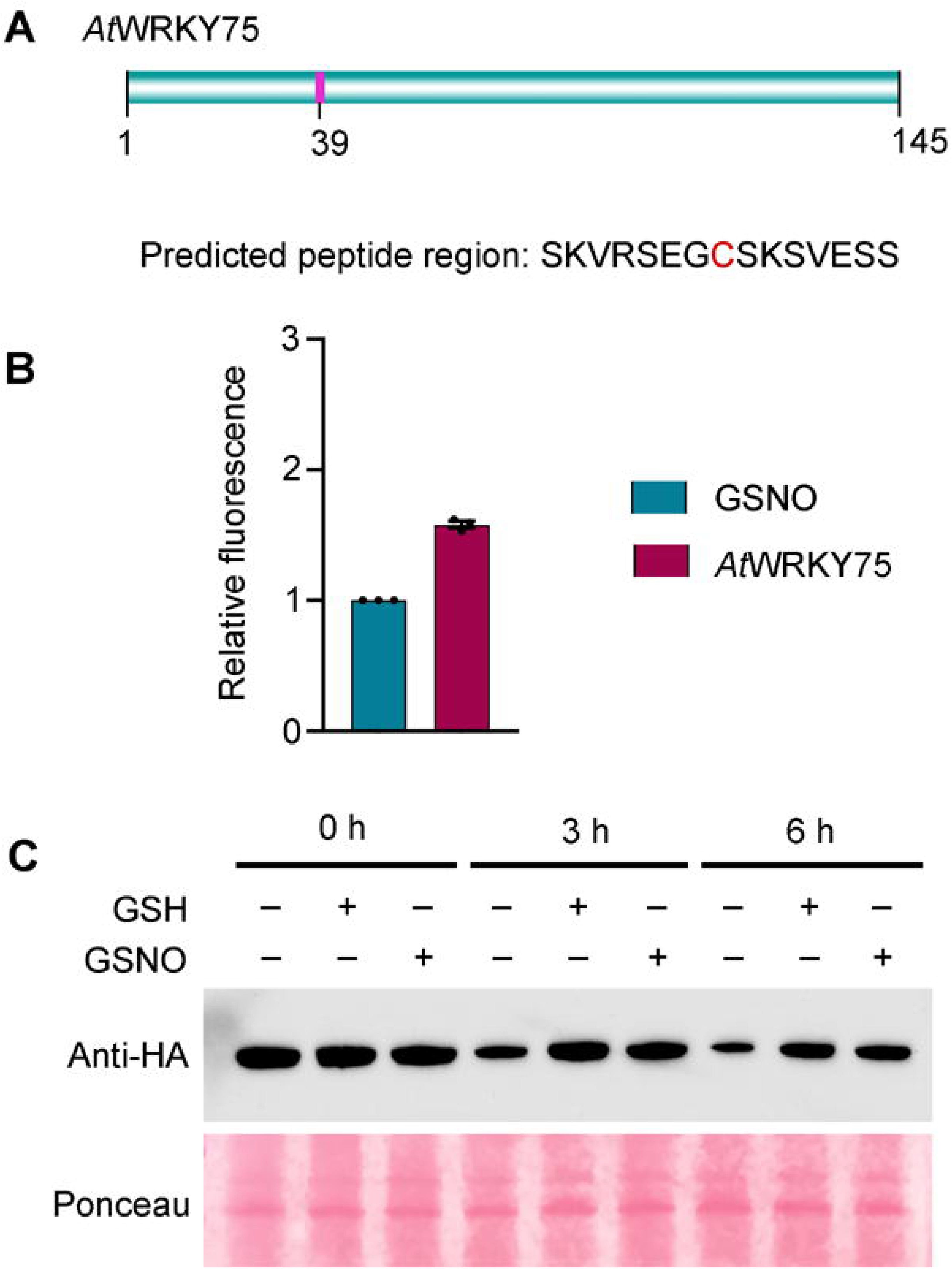
S-nitrosylation of WRKY 75 transcription factor (TF) (A) *In-silico* analysis for identification of the putative S-nitrosylation sites in WRKY75 TF amino acid sequence (B) DAN assay of S-nitrosylated Cys residue in wrky75 TF. Results were represented as mean±SEM of replicates in three independent experiments (C) protein stability assay of WRKY75 TF in response to exogenous GSNO treatment using anti-HA antibody. Ponceau staining was used as loading control. The protein stability assay was independently repeated thrice.

## Discussion

Over the last decade, accumulating evidence has revealed that GSH can regulate the transport and homeostasis of several nutrients, such as Fe and zinc, in plants (Shanmugam et al., 2011, 2015; Shee et al., 2021). A recent report has further demonstrated its role in regulating root hair growth under P deficiency via an indole-3-butyric acid-dependent pathway (Trujilo-Hernandez et al., 2020). However, no comprehensive investigation has been conducted to date to examine the precise mechanism of how GSH regulates P acquisition or reallocation in plants. In this study, we found that GSH regulates root and shoot P content in Arabidopsis, depending on Pi availability.

The two GSH-deficient mutants, *cad2-1* and *pad2-1* exhibited hypersensitivity to Pi deprivation with alterations in their RSA (Fig.1). Changes in RSA are an adaptive strategy for plants under Pi-limited conditions (Peret et al., 2014). Low Pi availability promoted lateral root growth over primary root growth by boosting lateral root density and length, and by limiting primary root growth through reduction in cell elongation (Williamson et al., 2001). The *pad2-1* mutant, which contains only 22% GSH, exhibited severe sensitivity to Pi deficiency with reduced lateral root density and rosette diameter compared to the WT. Another characteristic of P deficiency response is a change in pigment composition, which includes chlorophyll degradation and elevated anthocyanin accumulation (Marschner, 2012; Hernández and Munné-Bosch, 2015). With reduced photosynthetic rates, P-deficient plants have evolved to sustain photo-oxidative stress through anthocyanin accumulation (Hernández and Munné-Bosch, 2015). In this study, the GSH-deficient mutants displayed lower anthocyanin accumulation under Pi-deficient conditions than the WT, indicating enhanced sensitivity (Fig.1G). The chlorophyll loss under Pi deprivation was also maximal in the *pad2-1* mutant (Fig.1F). These observations resemble an earlier report where the *cad2-1* displayed increased sensitivity to Pi depletion (Trujilo-Hernandez et al., 2020), indicating that the optimum GSH level is essential for adaptation under Pi-limited conditions.

Plants have various P transport mechanisms comprising different high- and low-affinity transporters that allow P acquisition and distribution within the plant. Since source-sink relationships continually change with developmental stage and P availability, P transport in plants is an elaborate and complex process (Raghothama, 1999; Bucher et al., 2001). To analyse whether GSH can regulate Pi acquisition or distribution in plants, we measured Pi and total P content in GSH-deficient mutants and WT plants under Pi-sufficient and Pi-deficient conditions. Under Pi-sufficient conditions, both *cad2-1* and *pad2-1* mutants displayed higher P accumulation in shoots but lower P content in roots than the WT. The results, however, were reversed under Pi-deficient conditions (Fig. 2). Exogenously altering GSH content in Col-0 plants resulted in a similar P distribution pattern, confirming our previous observation (Fig. 3). These observations suggested a plausible role of GSH in modulating the P reallocation between roots and shoots based on the Pi availability status.

Mechanistically, Pi starvation elicited a characteristic defence response in WT roots. This was marked by a significant increase in the total GSH pool (Fig. 2). At the same time, the GSH:GSSG ratio declined across all genotypes, indicating that Pi deficiency shifts the root cellular redox milieu to a more oxidised state (Fig. 2). To determine whether this cascade results from general thiol-reduction potential changes or absolute GSSG accumulation, we tested the effects of the non-specific reducing agent DTT. DTT administration did not alter P/Pi content or *AtPHT1;5* transcript abundance under either Pi-sufficient or Pi-deficient conditions (Fig. 3, S1). Because DTT-mediated GSSG reduction did not suppress or mimic GSH fluctuation effects, these findings rule out a direct signalling role for GSSG or bulk redox shifts. This isolates reduced GSH and its downstream interaction with the GSNO module as the specific molecular signal governing Pi homeostasis.

This pattern is evident in the *pad2-1* mutant. Despite its extreme sensitivity to Pi starvation, *pad2-1* accumulates higher root Pi and total P than Col-0 under Pi-deficient conditions (Fig. 2). Analyses of shoot profiles and transcriptional data indicate this is not due to defective uptake, but rather a major failure in root-to-shoot P mobilisation. This failure results from GSH deficiency, which suppresses transcription of the key phosphate transporter *AtPHT1;5* (Fig. 3). Without this transporter, mutant roots lack the machinery to export Pi to the vascular tissue, causing remobilised or newly acquired P to be sequestered in the *pad2-1* root. Thus, our findings demonstrate that although Pi starvation changes root redox kinetics, a functional pool of reduced GSH is specifically required to activate *AtPHT1;5* expression, which in turn maintains source-to-sink Pi allocation. We interpret that this sequestration of Pi in roots may feedback to repress lateral root initiation in *pad2-1*. In contrast, *cad2*□*1*, which shows Pi retention in root similar to WT under Pi starvation, displays a more WT□like LR response. However, the differential sensitivity of LR formation to local versus systemic Pi signals, possible genotype-specific differences in hormone (e.g., auxin) or ROS signalling linked to GSH status needs to be investigated in future.

Several P transporters are known to regulate P homeostasis through P translocation and redistribution in plants (Mudge et al., 2002; Shin et al., 2004; Nagarajan et al., 2011; Lapis-Gaza et al., 2014). Our experiments identified that GSH regulates the expression of the *AtPHT1;5* under Pi-sufficient and Pi-deficient conditions (Fig.3). Among the PHT1 members, *AtPHT1;5* is expressed exclusively in phloem cells of older leaves, cotyledons, and flowers during P starvation (Mudge et al., 2002). Another genome-wide transcriptomic study reported *AtPHT1;5* expression in senescent Arabidopsis leaves (van der Graaff et al., 2006). Intriguingly, *At*PHT1;5 plays a pivotal role in transporting P from the source to the sink organs in response to developmental signals and P availability. During Pi starvation, the loss-of-function mutation in the *AtPHT1;5* gene resulted in decreased P allocation to shoots, whereas under Pi-replete conditions, the mutant exhibited higher shoot P content than the wild type but lower root P content (Nagarajan et al., 2011). Notably, the GSH-deficient mutants in our investigation displayed a strikingly similar P distribution pattern to that of the *pht1;5* mutant. These similarities in P distribution patterns between GSH-deficient mutants and the *pht1;5* mutant, along with the lack of response to exogenous GSH in *pht1;5* lines, indicate that *At*PHT1;5 mediates GSH-dependent source-to-sink P allocation. This suggests that GSH acts upstream of *At*PHT1;5, fine-tuning P transport in response to Pi availability. These findings strongly suggest that GSH not only regulates the expression of the *AtPHT1;5* gene but also integrates the source-to-sink P allocation mechanism via this transporter in accordance with P availability. However, it can also be considered that transcriptional regulation of a phloem□localized transporter like PHT1 can have amplified physiological consequences because small changes in transporter abundance or activity in phloem can disproportionately affect long distance P flux (Dai et al., 2022).

In Arabidopsis, GSH and NO combine to generate GSNO, a crucial signalling intermediate that modifies various nutrient transport genes (Kailasam et al., 2018; Koen et al., 2012; Shanmugam et al., 2015; Shee et al., 2022). It raised our curiosity about if GSNO is also involved in controlling the expression of the *AtPHT1;5* gene. Exogenous treatments with GSH, GSNO, or SNP result in greater gene induction under DP conditions, suggesting that the GSH-GSNO module plays positive role in *AtPHT1;5* gene regulation. This theory is further supported by the fact that the NO scavenger cPTIO eliminates the impact of GSH feeding and the GSH biosynthesis inhibitor BSO reverses the effect of the NO donor SNP (Fig. 5). In the NO-deficient *noa1* mutant, the expression of these transporter genes is markedly downregulated under Pi-deficient conditions but upregulated in the hyperaccumulating-NO *gsnor1-3* mutant. All of these outcomes indicate that GSH regulates *AtPHT1;5* gene regulation via the GSNO mediated pathway.

The next pertinent question is how the GSH-GSNO module regulates the *AtPHT1;5* expression. Since the module alters the relative transcript abundance of the transporter, transcriptional regulation is the most likely reason. Our experiments demonstrated that the *AtPHT1;5* promoter is inducible under Pi-deficient conditions. The promoter activity is also induced in response to exogenous GSH and GSNO treatment, indicating a GSH-GSNO module-mediated transcriptional regulation of the *AtPHT1;5* gene (Fig.6). However, being a metabolite, they cannot directly interact with the promoter to modulate its activity. Therefore, we were interested to identify the TF(s) involved in the process. Earlier reports suggested that GSH can activate different gene expression by modulating different TFs (Shee et al., 2022). For example, GSH can modulate genes in the ethylene biosynthetic pathway via the WRKY33 TF (Datta et al., 2015). In another study, we reported that under Fe-limited conditions, GSH can regulate the *NRAMP3/4* and *PIC1* genes by S-nitrosylating Fe-responsive bHLH TFs (Shee et al., 2022). Several TFs are reported to regulate the expression of different P starvation-induced genes. Previously, it was reported that the WRKY6 and WRKY42 TFs function as repressors of the P transporter, PHO1 (Chen et al., 2009). Under low P conditions, these TFs are degraded, allowing *PHO1* expression and subsequent regulation of P homeostasis. WRKY75 has long been known to regulate P acquisition and root development in plants (Devaiah et al., 2007). Another TF, WRKY45, can directly bind to the w-box of the *PHT1;1* promoter to activate its expression under Pi-limited conditions (Wang et al., 2014). Among the different MYB TFs, MYB2 and MYB62 play crucial roles in regulating the P starvation response. While MYB2 directly interacts with the promoter of miR399, a marker of P starvation response, MYB62 regulates the Pi starvation response via the gibberellic acid-mediated signalling pathway (Devaiah et al., 2009; Baek et al., 2013). In the present investigation, *At*WRKY42, *At*WRKY75, and *At*MYB2 were identified to interact with the *AtPHT;5* promoter. Among these three TFs, the *AtPHT1;5* expression was significantly lower in the *wrky7*5 mutant (Fig.7). This data corroborates the previous report where *AtPHT1;5* expression was diminished in the wrky75-RNAi lines under both P-sufficient and P-deficient conditions (Nagarajan et al., 2011). Moreover, *AtWRKY75* expression was significantly down-regulated in the GSH-deficient *pad2-1* mutant, which supports a previous report showing that GSH-treated Col-0 leaves displayed higher WRKY75 expression than the control condition (Sinha et al., 2014). We have further demonstrated that the *At*WRKY75 binds to the w-box motifs on the *AtPHT1;5* promoter, and its binding affinity increases in response to Pi starvation or GSH and GSNO treatment (Fig.7). These data delineate that GSH induces *AtPHT1;5* expressions under Pi-deficient conditions via the *At*WRKY75 TF.

It is generally known that S-nitrosylation of the TFs is involved in GSNO-mediated regulation (Darbani et al., 2013). Our previous study revealed that *At*bHLH38, *At*bHLH29, and *At*bHLH101 are stabilised by S-nitrosylation, and these TFs help regulate *AtNRAMP3/4* and *AtPIC1* expression in Fe-limited conditions (Shee et al., 2022). It is interesting to note that *At*WRKY75 protein has a single S-nitrosylation site, and GSNO treatment greatly improves its protein stability (Fig.8). These findings clearly imply that GSH regulates these proteins through S-nitrosylation, which similar to a previous study where the NO scavenger cPTIO was found to decrease *At*FIT1 stability (Meiser et al., 2011).

In summary, it can be concluded that in depleted phosphate conditions, GSH levels are elevated and are then converted to GSNO via S-nitrosylation. This GSNO causes S-nitrosylation of *At*WRKY75 TF. The S-nitrosylated *At*WRKY75 TF induces the transcription of *AtPHT1;5* gene which is directly involved in P allocation from source to sink during Pi starvation (Fig.9). Nevertheless, GSH likely influences P homeostasis at multiple levels like transcriptional, post translational, redox buffering, transporter activity, and metabolic redistribution. We have, therefore, concluded that while our data identify a key mechanistic pathway linking GSH-GSNO to *AtPHT1;5* induction via WRKY75, other GSH dependent processes may also contribute to the observed changes in P uptake and allocation, which can be explored in future.

**FIGURE 9.**
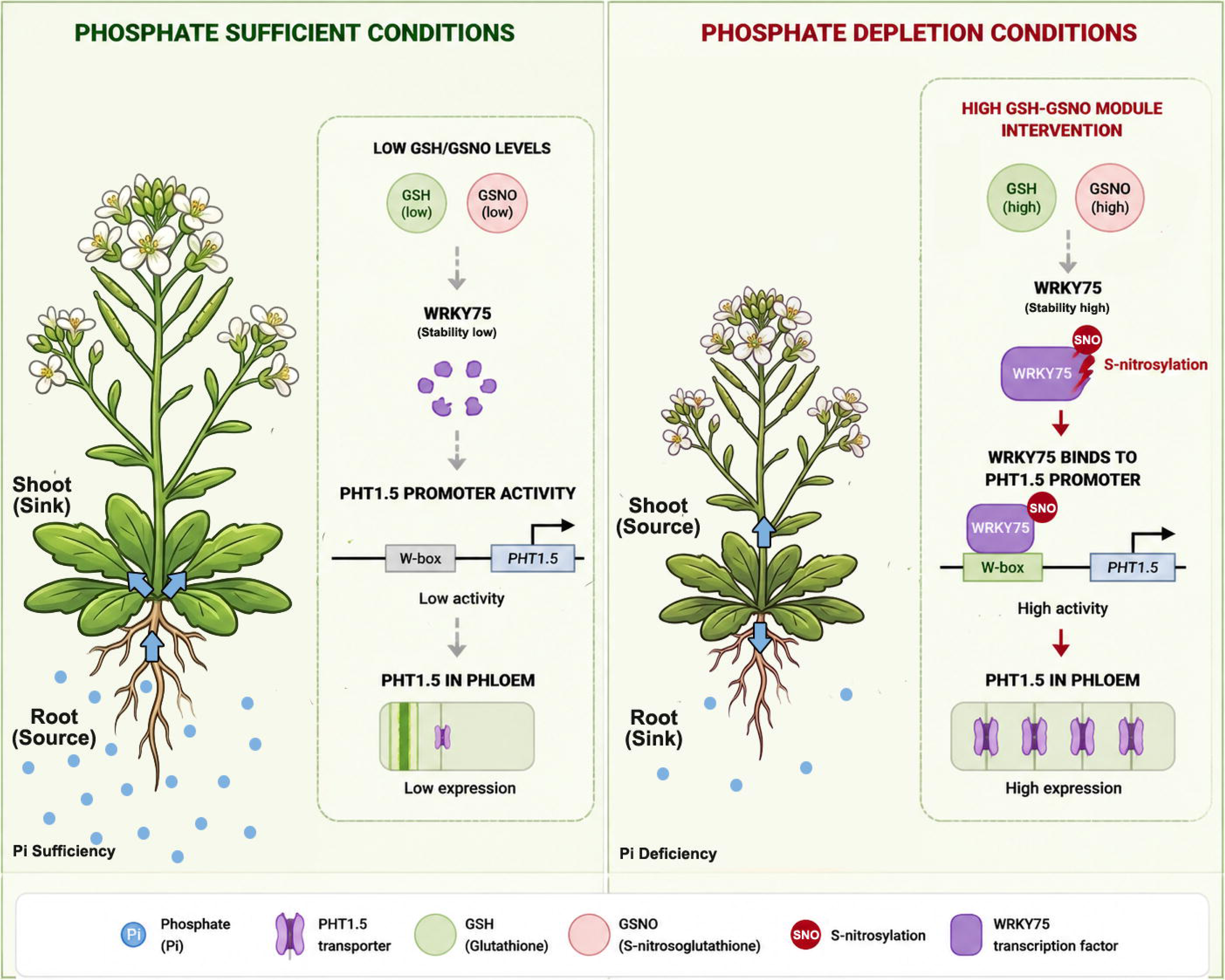
Proposed working model for GSH-GSNO mediated regulation of P translocation under Pi-limited condition. Pi deficiency leads to the accumulation of GSH which is converted to GSNO in cells and activates the P transporter, *AtPHT1;5* via S-nitrosylation of WRKY75 transcription factor. The accumulation of *At*PHT1;5 increases source to sink P distribution.

## Materials and Methods

### Plant growth and stress treatment

Seeds of *Arabidopsis thaliana* Columbia ecotype (Col-0), that served as WT, and two GSH-deficient mutants, *cad2-1* (N68137), and *pad2-1* (N3804), as well as *pht1;5* mutant (N574836) were procured from Nottingham Arabidopsis Stock Centre, UK. Seeds of *myb2* (SALK_043075), *wrky42* (SALK_049063), *wrky75* (CS121513), *noa1* (CS819466), *gsnor 1- 3* (CS66012) mutants were procured from the Arabidopsis Biological Resource Centre, USA. Seeds were surface-sterilised with 4% sodium hypochlorite and Tween 20, and germinated on standard MS medium. Plants were maintained in a growth chamber at 21±2°C and 60% relative humidity, with a 16 h light/8 h dark photoperiod, as standardised previously (Datta et al. 2015). For stress treatment, 7 d old seedlings were transferred to Pi-sufficient (1.25 mM KH_2_PO_4_, +P) or Pi-deficient (0 mM KH_2_PO_4_, supplemented with 0.625 mM K_2_SO_4_, -P) (Lu et al., 2017) media and maintained for 7 d. After the stress treatment, the plants were photographed, and morphological parameters, *viz*. primary root length, lateral root number, lateral root density, and rosette diameter, were analysed using ImageJ software.

### Estimation of Pi and total P content

Pi levels were determined using the ammonium molybdate method, as described by Lapis-Gaza et al. (2014). Root and shoot tissues were collected under Pi-sufficient or Pi-deficient conditions. Briefly, fresh tissues were homogenised in 1% (v/v) acetic acid and centrifuged at 12,000 g for 15 min at 4 . The supernatant was diluted and mixed with 0.35% (w/v) ammonium molybdate and 1.4% (w/v) ascorbic acid in 1 N sulphuric acid. The mixture was then incubated in the dark for 1 h. Absorbance was measured at 820 nm using a Double Beam UV-Vis Spectrophotometer (Hitachi). Pi concentration was determined from a standard curve prepared from KH_2_PO_4_ dilutions.

The total P content was measured using the ammonium blue method, as described by Nagarajan et al. (2011). Briefly, root and shoot tissues were dried in a hot air oven at 70 for 2 d. The dried samples were weighed and ashed at 500 for 3 h. The ash was dissolved in concentrated hydrochloric acid. The sample was diluted, then mixed with 0.35% (w/v) ammonium molybdate solution and incubated at 45 for 20 min. Optical density was measured at 820 nm.

### Estimation of chlorophyll and anthocyanin contents

200 mg tissue samples were homogenised in 80% acetone, followed by centrifugation at 5,000 g for 5 min. The total chlorophyll content was determined following Lichtenthaler (1987). Anthocyanin content was estimated following Matsui et al. (2004). Briefly, samples were homogenised in liquid nitrogen and resuspended in 5 volumes of 45% methanol and 5% acetic acid (v/v). Following centrifugation at 5,000 g for 5 min, absorbance was measured at 530 nm and 637 nm using a Double-Beam UV-Vis Spectrophotometer (Hitachi), and anthocyanin content was calculated.

### Estimation of total GSH content and determination of GSH:GSSG ratio

Reduced GSH content was estimated by the 5,5’-dithiobis-(2-nitrobenzoic acid) (DTNB) method as described by Anderson (1985). Briefly, 200 mg of tissue was homogenised in 5% sulphosalicylic acid and centrifuged at 12,000 g for 20 min. The supernatant was allowed to react with 50 µM DTNB in reaction buffer [0.1 M phosphate buffer (pH 7) and 0.5 mM EDTA] for 5 min, and the optical density was measured at 412 nm using a Double Beam UV-Vis Spectrophotometer (Hitachi). For the estimation of total GSH, 0.4 mM NADPH and 1 U glutathione reductase were added, and the mixture was incubated for 20 min. The optical density was measured at 412 nm. The amount of GSSG was determined by subtracting reduced GSH from total GSH.

### RNA extraction and quantitative-RT PCR analysis

The TRIzol method was used to isolate total RNA from the samples. The complementary DNA (cDNA) was prepared using the iScript™ cDNA Synthesis Kit (Bio-Rad) according to the manufacturer’s protocol. Quantitative PCR amplification was performed using the CFX96 Touch™ Real-Time PCR Detection System (Bio-Rad) using iTaq™ Universal SYBR^®^ Green Supermix (Bio-Rad) and gene-specific primers. *AtPHT1;4* was used as a marker gene for the Pi-deficient condition, and *AtActin*2 and *AtGAPDH* were used as reference genes.

### Chemical treatment of seedlings

15 d old seedlings were used for chemical treatments. The seedlings were placed on a petri plate containing filter paper wetted with half-strength MS medium with the appropriate concentration of the desired chemical(s) and sealed with parafilm. The seedlings were treated with freshly prepared 100 μM GSH (Sigma□Aldrich) or with 1 mM buthionine sulphoximine (BSO) (Sigma□Aldrich) for 72 h or with 5 mM dithiothreitol (DTT) (SRL) solution for 24 h. For GSNO feeding, the seedlings were treated with 250 μM GSNO (Sigma□Aldrich) for 72 h. The seedlings were also treated with 1 µM sodium nitroprusside (SNP) (SRL) and 0.5 mM 1,3□dihydroxy□4,4,5,5□tetramethyl□2□4 carboxyphenyl)tetrahydroimidazole (cPTIO) (Sigma-Aldrich) for 48 h, as previously standardised (Shee et al., 2022). Control seedlings were maintained in half□strength MS medium.

### Vector construction and raising of transgenic lines

Genomic DNA was extracted from Col-0 plants using the CTAB method. To generate various constructs, the 1500 bp sequence upstream of the +1 site of the *AtPHT1;5* gene was PCR-amplified with specific primers and cloned into the *pCAMBIA1304* vector between the *BglII* and *BamHI* restriction sites, producing the *pPHT1;5*::*GUS* construct. Coding sequences of *AtPHT1;5* and *AtWRKY75* were PCR-amplified from cDNA using gene-specific primers. *AtPHT1;5* was then cloned into the pBI121 vector between the *BamHI* and *SacI* sites to yield the *CaMV35S*::*PHT1;5* construct. The PCR-amplified *AtPHT1;5* promoter was further subcloned into the recombinant *CaMV35S*::*PHT1;5* vector (between *SphI* and *BamHI*) to generate the *pPHT1;5::PHT1;5* construct. Next, amplified *AtWRKY75* and *AtPHT1;5* were cloned into the *pVYCE* vector under the *CaMV35S* promoter between *SpeI/BamHI* and *SacI/BamHI* sites, respectively, creating *CaMV35S::WRKY75-HA* and *CaMV35S::PHT1;5-HA* constructs. The recombinant *pPHT1;5*::*PHT1;5* construct was used in *Agrobacterium*-mediated transformation of the *pht1;5* mutant to obtain complementation lines using floral dip method (Clough and Bent, 1998). *CaMV35S::WRKY75-HA* and *CaMV35S::PHT1;5-HA* constructs were similarly transformed into Col-0 plants to develop *AtWRKY75* and *AtPHT1;5* overexpression lines. Similarly, the *pPHT1;5*::*GUS* recombinant construct was transformed into Col-0 and *wrky75* mutant lines, while vector control lines containing the *CaMV35S::GUS* construct were generated in both Col-0 and *wrky75* backgrounds. Putative transformants were selected via antibiotic screening (kanamycin for *pPHT1;5*::*GUS* and *pPHT1;5*::*PHT1;5,* and hygromycin for *CaMV35S::WRKY75-HA* and *CaMV35S::GUS*) and confirmed by selectable marker-specific PCR. Selected plants were grown to the T_2_ generation in a growth chamber as described above.

### Histochemical GUS assay

The 7 d old seedlings from transgenic lines (T_2_ generation) harbouring *pPHT1;5::GUS* and *CaMV35S::GUS* constructs (in the Col-0 background) were maintained under either +P, -P, +GSH, or +GSNO conditions as described above. Following treatment, samples were infiltrated with GUS staining solution (0.5 mg mL^−1^ 5-bromo-4-chloro-3-indolylβ-D-glucuronic acid, 0.5 mM potassium ferrocyanide, 0.5 mM potassium ferricyanide, 0.1% [v/v] Triton X-100, 100 mM phosphate buffer, pH 7.0, and 10 mM EDTA), and a histochemical GUS assay was performed as described by Jefferson et al. (1987). Finally, samples were washed with 70% alcohol and photographed.

### In silico promoter analysis and prediction of S-nitrosylation sites

The presence of the w-box motif was identified in the promoter region of *AtPHT1;5* using PlantPan 3.0 (https://plantpan.itps.ncku.edu.tw/promoter.php) and MEME suite 5.3.3 (https://memesuite.org/). To identify the presence of S□nitrosylated cysteine residues in the targeted TFs, the protein sequences were retrieved from the NCBI database, and the sequences were analysed with the GPS□SNO 1.0 software (Xue et al., 2010).

### Y1H analysis

The PCR-amplified promoter sequence of the *AtPHT1;5* gene was cloned into the *pAbAi* vector (Takara) between the *SacI* and *KpnI* RE sites, following the manufacturer’s protocol. The cds of *AtWRKY6, AtWRKY42, AtWRKY45, AtWRKY75, AtMYB2,* and *AtMYB62* genes were amplified by PCR using gene-specific primers and cloned into *pGADT7-Rec* vector (Takara). The recombinant constructs were transformed into the Y1H gold yeast strain (Takara). The transformed strains were grown on SD/-Leu and SD/-Leu/Aureobasidin A media (Takara), and Y1H analysis was performed using Matchmaker^®^ Gold Yeast One-hybrid screening system (Takara) according to the manufacturer’s protocol. The interaction between p53-*pAbAi* was used as a positive control, and *pAbAi* alone was considered a negative control.

### Dual luciferase transactivation assay

The promoter region of *AtPHT1;5* was cloned into the *pGreen-LUC008* vector between *SpeI* and *KpnI* RE sites to drive the expression of the firefly luciferase gene. The construct constitutively expressing the Renilla luciferase (*REN*) gene and harbouring the *pPHT1;5::LUC* cassette was used as reporter. The *CaMV35S::WRKY75-HA* construct was considered as an effector. *Agrobacterium* strains harbouring the recombinant constructs (effector and reporter) were mixed in a 1:1 (v/v) ratio and used for *Agrobacterium*-mediated infiltration of *Nicotiana tabacum* leaves as standardized before (Sahid et al., 2020). For the negative control, an *Agrobacterium* strain harbouring the effector construct and an empty *pGreen-LUC008* vector were mixed in a 1:1 (v/v) ratio and used for infiltration (Zhang et al., 2015). After incubation for 48 h in the dark, infiltrated leaves were harvested, and total protein was isolated. 1 µg protein was used for a dual luciferase transactivation assay using Dual-Luciferase^®^ Reporter Assay System (Promega) following the manufacturer’s protocol. Luminescence was measured with the Varioskan LUX luminometer (Thermo Scientific), and relative luciferase activity was calculated as the LUC/REN ratio.

### CHIP-qPCR analysis

The *CaMV35S::WRKY75-HA* harbouring transgenic line was used for the ChIP assay following Cheng et al. (2020) with slight modifications. Briefly, seedlings were maintained under +P or -P conditions or treated with exogenous GSH or GSNO as described above. 800 mg tissue from each sample was harvested, washed with milliQ water, cross-linked with 1% (v/v) formaldehyde solution under vacuum for 10 min, and homogenised in liquid nitrogen. The chromatin complex was isolated and sheared by sonication (Hielscher Ultrasonic processor, UP200S). The complex was then incubated with 2 µg of anti-HA antibody (Sigma). The precipitated DNA was recovered, and ChIP qPCR was performed using EpiQuik™ Chromatin Immunoprecipitation (ChIP) Kit (Epigentek). For the negative control, non-immune IgG antibody was used. Real-time PCR was performed in triplicate using a CFX96 Touch™ Real-Time PCR Detection System (Bio-Rad) with iTaq™ Universal SYBR^®^ Green Supermix (Bio-Rad) and two sets of w-box-specific primers for the *AtPHT1;5* promoter. Chromatin without antibody treatment was used as input DNA. The % input was calculated from Ct values.

*In vivo 2,3*□*diaminonapthalene (DAN) assay of S*□*nitrosylated proteins*

The *CaMV35S::WRKY75-HA* harbouring transgenic line was used for this assay. 2 g of tissue sample was homogenized in lysis buffer (50 mM Tris, pH 7.5, 150 mM sodium chloride, 10 mM magnesium chloride, 1% glycerol, 0.5% Triton X□100, and 1 mM phenylmethylsulfonyl fluoride as protease inhibitor). Following centrifugations at 13,000 g for 10 minutes at 4°C twice, the sample was filtered through a 0.22 μm PVDF membrane to collect the supernatant. After adding 80 µl of anti-HA agarose beads, the supernatant was incubated for 2 h at 4°C. Next, 400 µM GSNO was added to it and incubated for 30 min in the dark. The sample was then rinsed with phosphate-buffered saline and incubated for 30 min in the dark with 300 µl of 200 µM DAN and 200 µM mercuric chloride. To stop the reaction, 2.8 M NaOH was added. Using an excitation wavelength of 365 nm and an emission wavelength of 450 nm, the Varioskan™ LUX multimode microplate reader (Thermo Scientific) was used to measure the fluorescence signal of the 2,3-naphthyltriazole (NAT) compound in the sample. The Bradford method was then applied to the solution to determine the protein concentration. The relative fluorescence intensity of the NAT molecule per micromolar of protein was used to calculate the DAN-NAT conversion rate of the sample.

### Protein stability assay

The *CaMV35S::WRKY75-HA* harbouring transgenic line was used for the protein stability assay. Briefly, seedlings were treated with exogenous GSH or GSNO as described above. After 72 h, the seedlings were treated with 200 µM cycloheximide, and total protein was isolated at 0, 3, and 6 h from GSNO- and GSH-treated and non-treated plants. Proteins were isolated using lysis buffer (20 mM Tris□Cl, pH 7.5, 150 mM sodium chloride, 0.5% Triton□X□100, and protease inhibitor) at 4°C, then centrifuged at 13,000 g for 30 min at 4°C. The supernatant was collected and quantified by the Bradford method, and immunoblot analysis was performed using an anti□HA antibody (1:1000 dilution, Sigma).

### Statistical analysis

Statistical analyses were performed using GraphPad Prism version 8.3.0 (GraphPad Software, San Diego, California, USA). Variations in morphological and biochemical parameters, as well as relative transcript abundance, were evaluated using one-way or two-way ANOVA, followed by Dunnett’s multiple-comparison test. Statistical significance between two data sets was determined at P≤0.05. Details regarding statistical tests, replicates, sample sizes, and P-values are provided in the respective figure legends. Data are presented as mean ± standard error of the mean (SEM).

## Conflict of interest

Authors declare no conflict of interest.

## Author Contributions

RD and SP conceived and designed the research plan; RS performed most of the experiments; DS performed *in silico* analysis and Y1H analysis, TS and RS performed DAN assay and protein stability assay, SS generated the constructs, RD and SP analyzed the data. RS, RD and SP wrote and finalized the manuscript.

## Acknowledgement

We thank the DST-FIST and DBT-BUILDER supported central instrumentation facilities of the Department of Botany, and Department of Biochemistry, University of Calcutta and Department of Botany, Dr A. P. J. Abdul Kalam Government College. RS acknowledges the University Research Fellowship, University of Calcutta for her fellowship.

## Supplemental information

**Figure S1** Chemical treatment of Col-0 seedlings in Pi-deficient condition

**Figure S2** Response of *pad2-1* mutant in response to exogenous GSH feeding.

**Figure S3** Expression of *AtPHT1;5* gene in mutant and *AtPHT1;5* mutant complementation lines after GSH and BSO treatment

**Figure S4** Expression of *AtPHT1;4* gene in mutant and AtPHT1;5 mutant complementation lines in response to Pi-deficient condition.

**Figure S5:** Position of the w-box motifs on the *AtPHT1;5* promoter.

**Figure S6:** Prediction of S-nitrosylation sites in the amino acid of WRKY75

**TABLE S1** List of primers used.

## Notes

**Funding:** This work has been supported by the Science and Engineering Research Board, Government of India [ECR/2017/000231] and Start up Grant, University Grant Commission, Government of India, India [F30-363/2017(BSR)]

### Competing Interest Statement

The authors have declared no competing interest.

### Summary of Updates

Section on Result, Discussion and Materials & Methods have been revised to clarify the role of glutathione and s-nitrosoglutathione to regulate AtPHT1.5 gene in Arabidopsis. All Figures have been revised, author list was revised, supplemental files updated

