## Supplemental Figures for "Glutathione-mediated S-Nitrosylation of WRKY75 modulates *PHT1;5* expression to orchestrate phosphate homeostasis in *Arabidopsis*"

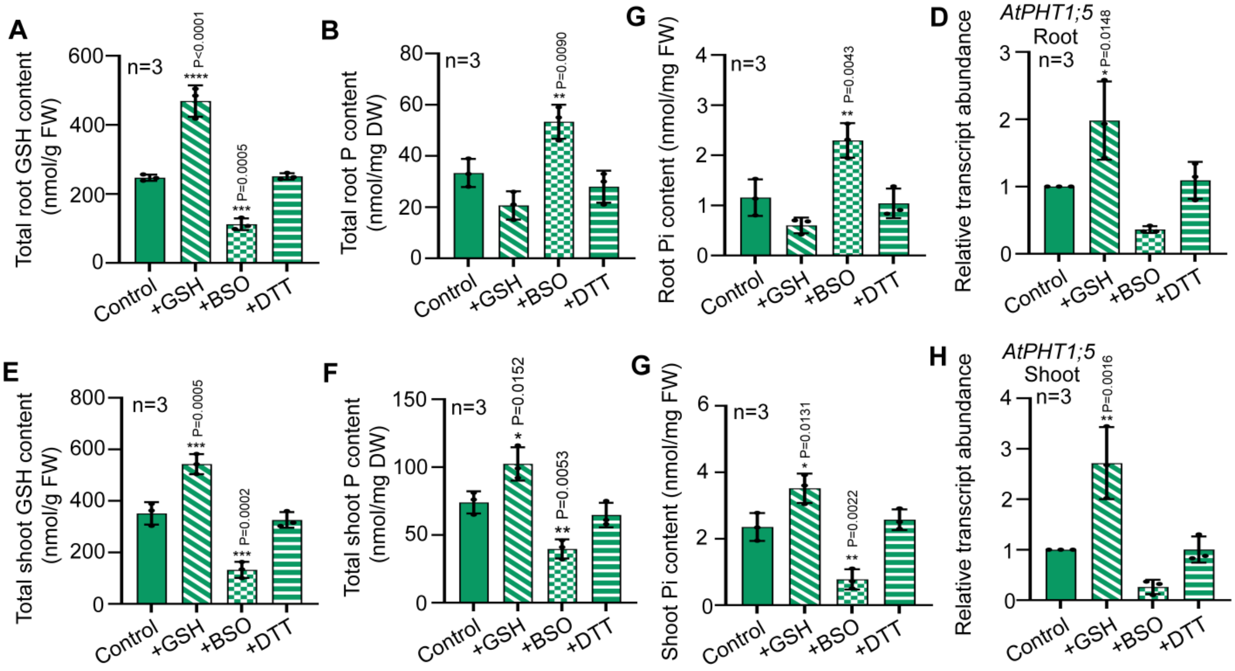


**Figure S1:** Chemical treatment of Col-0 seedlings in Pi-deficient condition. 7 d old MS grown seedlings of Col-0 plants were exposed to -P conditions for 7 d and treated with 100 μM GSH for 72 h, or 1 mM BSO for 72 h, or 5 mM DTT solutions for 24 h. Control plants were maintained in half strength MS medium for the entire duration and analyzed. (A) Total GSH content in roots (B) total root P content (C) root Pi content (D) relative transcript abundance of AtPHT1;5 in roots, (E) total GSH content in shoots (F) toal shoot P content (G) shoot Pi content, (H) relative transcript abundance of AtPHT1;5 gene in shoots. The experiment was independently repeated thrice and results were represented as mean±SEM (n=3). Statistical differences between the treatments were analyzed by one-way ANOVA followed by Dunnett's multiple comparison test and statistical significances were denoted by asterisks in the respective panels.

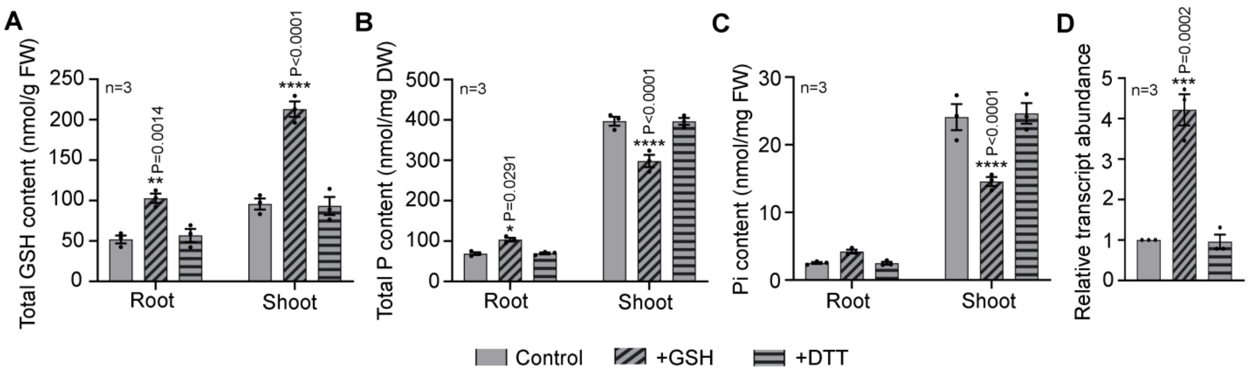


**FIGURE S2 Response of *pad2-1* mutant in response to exogenous GSH feeding.** 14 d old MS grown *pad2-1* plants were treated with 100 µM GSH for 72 h, or 1 mM BSO for 72 h, or 5 mM DTT for 24 h solutions and analyzed. Control plants were maintained in half strength MS medium for the entire duration. (A) Total GSH content, (B) total P content, (C) Pi content, and (D) relative transcript abundance of AtPHT1;5 gene in *pad2-1* seedling. The experiment was independently repeated thrice and results were represented as mean±SEM (n=3). Statistical differences between the treatments were analyzed by two-way ANOVA (A-C) or one-way ANOVA (D) followed by Dunnett's multiple comparison test and statistical significances were denoted by asterisks in the respective panels.

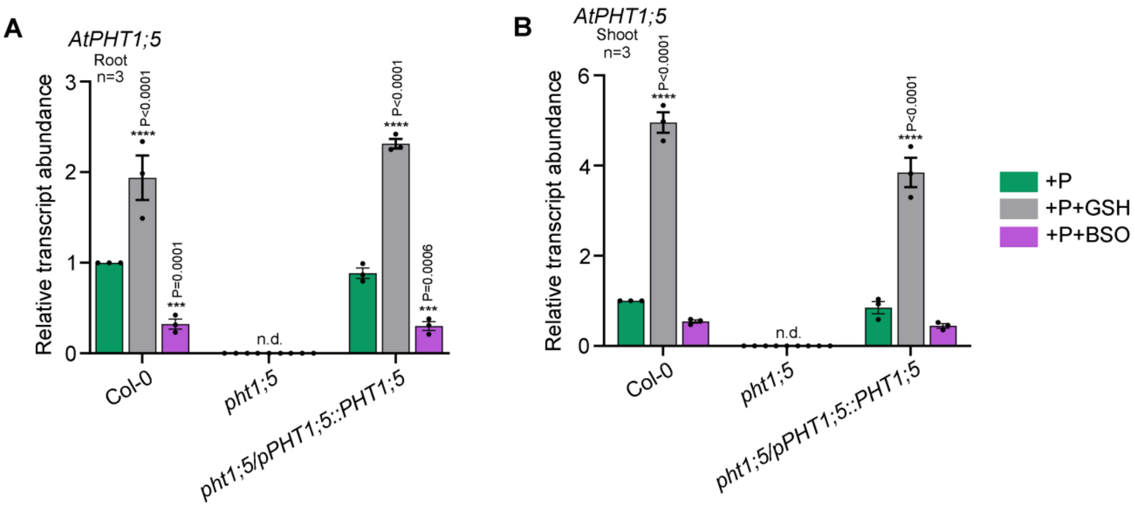


**Figure S3:** Expression of *AtPHT1;5* gene in *Atpht1;5* mutant and *Atpht1;5* mutant complementation lines after GSH and BSO treatment 14 d old MS grown *Atpht1;5* mutants and complementation plants were treated with 100 µM GSH, or 1 mM BSO for 72h and analyzed. (A) Relative transcript abundance of AtPHT1;5 gene in roots of Col 0, pht1;5 mutant and complementation lines (B) relative transcript abundance of AtPHT1;5 gene in roots of Col 0, pht1;5 mutant and complementation lines. The experiment was independently repeated thrice and results were represented as mean±SEM (n=3). Statistical differences between the treatments were analyzed by two-way ANOVA followed by Tukey’s multiple comparison test and statistical significances were denoted by asterisks in the respective panels.

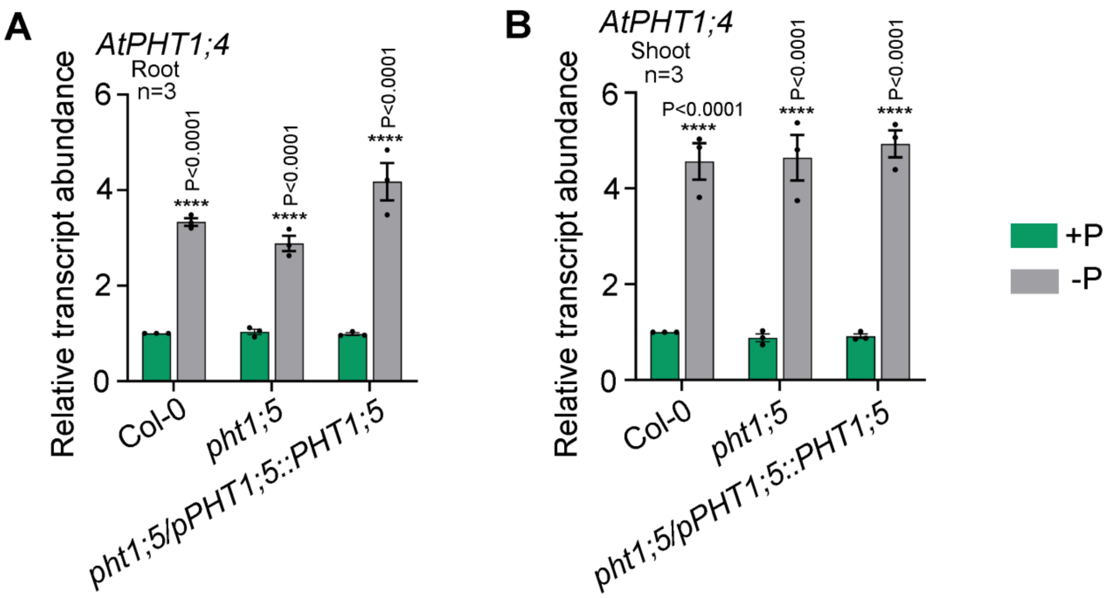


**Figure S4:** Expression of *AtPHT1;4* gene in Atpht1;5 mutant and Atpht1;5 mutant complementation lines in response to Pi-deficient condition**.** 7 d old MS grown seedlings of Col-0, *pht1;5* mutant and complementation plants were exposed to -P conditions for 7 d and analyzed. (A) relative transcript abundance of *AtPHT1;4* gene in roots (B) relative transcript abundance of AtPHT1;4 gene in shoot. The experiment was independently repeated thrice and results were represented as mean±SEM (n=3). Statistical differences between the treatments were analyzed by two-way ANOVA followed by Sidak’s multiple comparison test and statistical significances were denoted by asterisks in the respective panels.

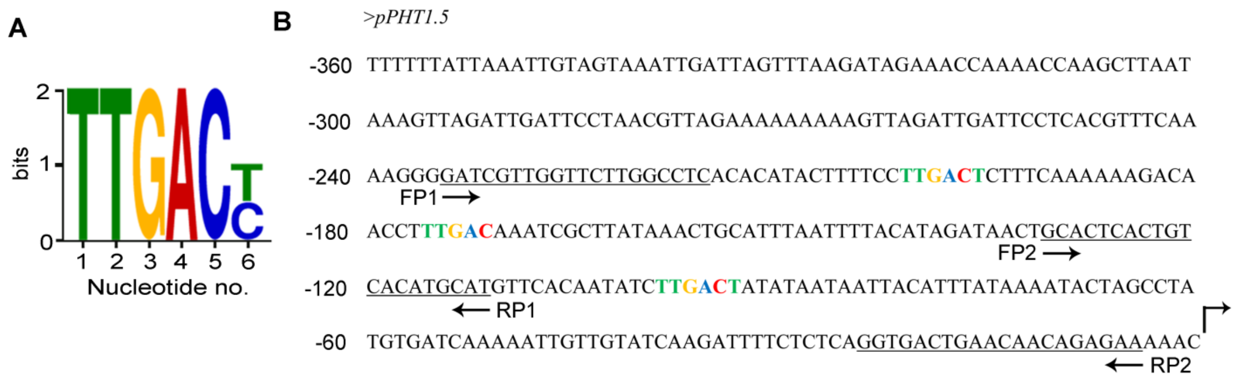


**FIGURE S5 Position of the w-box motifs on the *AtPHT1;5* promoter.** (A) w-box motif, and (B) position of w-box motif and primers used in ChIP-qPCR analysis on the promoter sequence.

**Predicted Site**

| Position | Peptide | Score | Cutoff | Cluster |
|---|---|---|---|---|
| 39 | SKVRSEGCSKSVESS | 2.75 | 2.443 | Cluster B |

**Figure S6: Prediction of S-nitrosylation sites in the amino acid of WRKY75**.
