## Supplemental table for "Glutathione-mediated S-Nitrosylation of WRKY75 modulates *PHT1;5* expression to orchestrate phosphate homeostasis in *Arabidopsis*"

**TABLE S1** List of primers used

| **Gene name** | **Loci number** | **Primers** |
|---|---|---|
| *AtPHT1;5* | AT2G32830 | Forward:5’ GGATCCCAACAGAGAAAAACACATGGCGAA 3’ |
| | | Reverse: 5’ GAGCTCTCAAACCGGGACTTTTCTACC 3’ |
| *AtPHT1;5* (partial) | AT2G32830 | Forward:5’CATCGCCTATTACAGCAGCA 3’ |
| | | Reverse:5’CGTCGCATTAGGTCCAAAAT 3’ |
| *AtPHT1;5* promoter (LUC) | AT2G32830 | Forward: 5’ GGTACCATACGTCGTGTAGCTTGAAT 3’ |
| | | Reverse: 5’ ACTAGTTTTTCTCTGTTGTTCAGTCAC 3’ |
| *AtPHT1;5* promoter | AT2G32830 | Forward: 5’GGATCCCGTTGATACTACAAAGATTCATG 3’ |
| | | Reverse: 5’CCATGGTTTCTCTGTTCAGTCACCT 3’ |
| *AtActin* | At3G18780 | Forward: 5’GCACCCTGTTCTTCTTACCG 3’ |
| | | Reverse: 5’AACCCTCGTAGATTGGCACA 3’ |
| *AtPHT1;5* promoter (Y1H) | AT2G32830 | Forward: 5’ GAGCTCCGTCGTGTAGCTTGAATATATTC 3’ |
| | | Reverse: 5’ GGTACCTTTTTCTCTGTTGTTCAGTCACG 3’ |
| *AtWRKY75* | AT5G13080 | Forward: 5’ACTAGTATGGAGGGATATGATAATGGGT 3’ |
| | | Reverse: 5’GGATCCCTAGAAAGAAGAGTAGATTTGCATT 3’ |
| *AtWRKY75* promoter | AT5G13080 | Forward: 5’ GGATCCACCGTATCTCACCACAACGA 3’ |
| | | Reverse: 5’ CCATGGTATAACAACGAGACAGACCG 3’ |
| *AtWRKY75* (Y1H) | AT5G13080 | Forward: 5’ ATGGAGGGATATGATAATGGGT 3’ |
| | | Reverse: 5’ CTAGAAAGAAGAGTAGATTTGCATT 3’ |
| *AtWRKY42* (Y1H) | AT4G04450 | Forward: 5’ATGTTTCGTTTTCCGGTAAG 3’ |
| | | Reverse: 5’ ATTGCCTATTGTCAACGTTG 3’ |
| *AtWRKY45* (Y1H) | AT3G01970 | Forward: 5’ ATGGAGGATAGGAGGTGTGA 3’ |
| | | Reverse: 5’ CTTCAAGCAAAAGGGAGGGA 3’ |
| *AtMYB2* (Y1H) | AT2G47190 | Forward:5’ ATG GAAGATTACGAGCGAAT 3’ |
| | | Reverse: 5’ TACGAATACGATGTCGTATC 3’ |
| *AtMYB62* | AT1G68320 | Forward: 5’ GTATGGAAAATTCGATGAAG 3’ |

| | | |
|---|---|---|
| (Y1H) | | Reverse: 5’ TACTCCCTAAACTGCCAAAT 3’ |
| *AtPHT1;1* (partial) | AT5G43350 | Forward: 5’TTGCCTTCCCTTACAACCAC 3’ |
| | | Reverse: 5’GCTAACCTCAGCCTCACCAG 3’ |
| *AtPHT1;2* (partial) | AT5G43370 | Forward: 5’GTGGAACCCTTTCTGGTCAA 3’ |
| | | Reverse: 5’AACTTGAGGAGGCGTTGAGA 3’ |
| *AtPHT1;7* (partial) | AT3G54700 | Forward: 5’ GGCATTGGTGGTGATTATCC 3’ |
| | | Reverse:5’ GTTTGAAGCCGCTAGTTTCG 3’ |
| *AtPHT1;8* (partial) | AT1G20860 | Forward: 5’CCTGCTGCATTGACGTTCTA 3’ |
| | | Reverse: 5’TGCGTCGTAGACGTTTTCTG 3’ |
| *AtPHT1;9* (partial) | AT1G76430 | Forward: 5’ GCCACGATTATGTCGGAGTT 3’ |
| | | Reverse: 5’ CATGTCTTTTGCTGCTTGGA 3’ |
| *AtPHT1;6* (partial) | AT5G43340 | Forward: 5’CTCCGGTATGGGTTTCTTCA 3’ |
| | | Reverse: 5’ AGGATAGTCACCACCGATGC 3’ |
| *AtPHT1;3* (partial) | AT5G43360 | Forward: 5’AAACTCGGACGGAAAAAGGT 3’ |
| | | Reverse: 5’ GCTTGAGGAGGCGTTGATAG 3’ |
| *AtPHT1;4* (partial) | AT2G38940 | Forward: 5’CTACCCTTTATCCGCAACCA 3’ |
| | | Reverse: 5’CAACCAAAGCCGTGTACCTT 3’ |
| *AtPHO1* (partial) | AT3G23430 | Forward: 5’ TGGTTCTCCGGAACAAGAAC 3’ |
| | | Reverse: 5’ GTCCCTGTCAAGGAACGGTA 3’ |
| *AtPHO2* (partial) | AT2G33770 | Forward: 5’ AGAAACCATTGGCAAAATCG 3’ |
| | | Reverse: 5’ GGTGACAGAGACACGCTCAA 3’ |
| *AtPHL1* (partial) | AT5G29000 | Forward: 5’ CATGAAAACGAGCGTTGAGA 3’ |
| | | Reverse: 5’ CACGTTTCCTTGATGCTGAA 3’ |
| *AtPHR1* (partial) | AT4G28610 | Forward: 5’ CGTTGATCTGGCAAACAGAA 3’ |
| | | Reverse: 5’ CATCTTTACAACGCCGGATT 3’ |
| *AtPHR2* (partial) | AT2G47590 | Forward: 5’ GACTAGCCAGGAGGGGAAAC 3’ |
| | | Reverse: 5’ CTTGGAGTTGTGGAGGTGGT 3’ |
| *AtPDR2* (partial) | AT5G23630 | Forward: 5’ CTCAGAATGGGGAAGGATCA 3’ |
| | | Reverse: 5’ TGTGGCGAGACAGTTCAGAC 3’ |
| *AtWRKY75* | AT5G13080 | Forward: 5’ ATTCGAAACCAGAGCTGCAT 3’ |

| | | |
|---|---|---|
| (partial) | | Reverse: 5’ ACGACCACTTCTTGGTCCAC 3’ |
| *AtWRKY42* (partial) | AT4G04450 | Forward: 5’ CACAATGGCTGTTGGATGTC 3’ |
| | | Reverse: 5’ AATGGTGCAGACGCTGAGAT3’ |
| *AtMYB2* (partial) | AT2G47190 | Forward: 5’ CTCTGGGCTAAAGCGAACTG 3’ |
| | | Reverse: 5’ TTCCACTAATCTCGGCATCC 3’ |
| *AtWRKY75* ChIP1 | AT5G13080 | Forward: 5’ GATCGTTGGTTCTTGGCCTC 3’ |
| | | Reverse: 5’ ATGCATGTGACAGTGAGTGC 3’ |
| *AtWRKY75* ChIP 2 | AT5G13080 | Forward: 5’ GCACTCACTGTCACATGCAT 3’ |
| | | Reverse: 5’ TTCTCTGTTGTTCAGTCACC 3’ |
